# Outer membrane utilisomes mediate oligosaccharide uptake in gut Bacteroidetes

**DOI:** 10.1101/2022.08.15.503959

**Authors:** Joshua B. R. White, Augustinas Silale, Matthew Feasey, Tiaan Heunis, Yiling Zhu, Hong Zheng, Akshada Gajbhiye, Susan Firbank, Arnaud Baslé, Matthias Trost, David N. Bolam, Neil A. Ranson, Bert van den Berg

## Abstract

Bacteroidetes are abundant members of the human microbiota, with species occupying the distal gut capable of utilising a myriad of diet- and host-derived glycans. Transport of glycans across the outer membrane (OM) of these bacteria is mediated by SusCD protein complexes, comprising a membrane-embedded barrel and a lipoprotein lid, that are thought to operate via a ‘pedal-bin’ mechanism in which the lids open and close to facilitate substrate binding. However, additional cell surface-exposed lipoproteins, namely surface glycan binding proteins and glycoside hydrolases, play critical roles in the capture and processing of large glycan chains into transport-competent substrates. Despite constituting a crucial mechanism of nutrient acquisition by our colonic microbiota, the interactions between these components in the OM are poorly understood. Here we show that for the levan and dextran utilisation systems of *Bacteroides thetaiotaomicron,* the additional OM components assemble on the core SusCD transporter, forming stable glycan utilising machines which we term ‘utilisomes’. Single particle electron cryogenic electron microscopy (cryo-EM) structures in the absence and presence of substrate reveal concerted conformational changes that rationalise the role of each component for efficient nutrient capture, as well as providing a direct demonstration of the pedal bin mechanism of substrate capture in the intact utilisome.

## Introduction

The human large intestine is home to a diverse microbial community, known collectively as the gut microbiome, which plays an essential role in human health^1^. Within the large intestine, complex dietary glycans which are inaccessible to the enzymes of the human digestive tract are the primary nutrient source, and the utilisation of these complex sugars by gut microorganisms is essential for their survival^2, 3^. Crucially, this nutrient supply is also integral to mutualism between host and bacteria, via the generation of short-chain fatty acids that are accessible to the host. The availability of such metabolites is associated with normal gastrointestinal physiology and systemic health benefits^4, 5^. The distal gut microbiome is dominated by two bacterial phyla: the Gram-negative Bacteroidetes and the Gram-positive Firmicutes^6^. The Bacteroidetes employ a common strategy for glycan utilisation. The machinery required for the uptake, processing and metabolism of specific glycans is encoded in co-regulated gene clusters known as polysaccharide utilisation loci (PULs)^7^.

The OM of Gram-negative bacteria presents a formidable barrier to the uptake of large nutrients^8^. Translocation of complex glycans across the Bacteroidetes OM is dependent on the SusCD core components of a PUL, comprising a TonB-dependent active transporter (SusC) and an associated lipoprotein (SusD)^7, 9^. Previous structural studies have shown that these SusCD complexes exist as SusC_2_D_2_ tetramers^10–12^, creating a core transportation unit with a twin barrel structure, where each barrel is associated with its own SusD component that caps the extracellular face of the transporter barrel. Recent work has shown that these SusCD-like transporters utilise a ‘pedal bin’ mechanism of nutrient uptake, where SusD is able to undergo large, hinge-like movements that alternatively expose or occlude a binding site for the nutrient within the lumen of the SusC barrel, facilitating substrate capture^12^.

Although most glycan breakdown occurs within the bacterial cell, the initial binding and processing of long glycan chains occurs extracellularly. To achieve this, an authentic PUL minimally encodes, in addition to at least one SusCD pair, a surface-located, endo-acting glycanase (most commonly glycoside hydrolases; GH) and one or more OM surface glycan binding proteins (SGBPs)^7, 9^. The model gut symbiont *B. thetaiotaomicron* (*B. theta*) has 88 predicted PULs comprising almost 20% of its genome, but the substrate specificity is known for only ∼20 of these^13^. The large number of PULs, each likely dedicated to the acquisition of a specific glycan, demonstrates the importance of complex glycans to bacterial survival in the distal gut.

One of the best-characterised PULs is that for the utilisation of levan, a plant and bacterial-derived fructan polysaccharide, which consists of β2,6-linked fructose units with occasional β2,1 branches^14, 15^. We previously characterised the binding and uptake of β2,6 fructo-oligosaccharides (FOS) by the core SusCD levan transporter^10, 12^. However, the important steps preceding levan capture and transport by SusCD remain unclear. Generation of transport-competent FOS from levan is achieved by the OM lipoprotein Bt1760 (a GH32-family levan endo-glycanase)^14, 16^, while a second OM lipoprotein, Bt1761 (the surface glycan binding protein; SGBP) has a putative role in recruitment of levan to the cell surface^17^. These two lipoproteins will henceforth be indicated as the levanase and the levan SGBP respectively, and collectively as the “additional lipoproteins”.

A key unresolved question in the field is whether and how the additional lipoproteins associate with their cognate SusCD transporters to drive oligosaccharide uptake. Some information is available for the archetypal starch utilisation system, which encodes two SGBPs (SusE and SusF) in addition to the SusCD transporter and a surface amylase (SusG)^18, 19^. Seminal studies offered evidence in support of SusCD-SGBP complex formation^19^. Since then, a more dynamic picture of the starch utilisation system, involving all 5 OM components, has emerged. Those studies indicate transient, substrate-induced complex formation^20^,and describe a role for the SGBPs SusE and SusF as immobile starch binding centres around which the SusCD transporter and the SusG amylase can assemble^21^. In contrast, the Bt2263^SusC^- Bt2264^SusD^ complex from an uncharacterised PUL in *B. theta* form a stable complex with two lipoproteins of unknown function (Bt2261 and Bt2262)^10^.Thus, broadly speaking, two opposing models exist: (1) The dynamic assembly model where free lipoproteins and the SusC transporter transiently assemble, possibly in response to the presence of substrate, and (2) The stable complex model where the additional lipoprotein components assemble on the core SusCD transporter.

Here we demonstrate, using proteomics and single particle cryo-EM, that all four OM components of the levan PUL from *B. theta* exist in a stable, octameric complex that we term a ‘utilisome’ in keeping with the “PUL” acronym. We also show that the four OM components of the *B. theta* dextran (a bacterial α-glucan) PUL adopt a similar architecture, suggesting that utilisomes are a generic feature of glycan utilisation in gut *Bacteroides*. Cryo-EM structures of both levan and dextran utilisomes in the absence of their cognate substrates suggest that the auxiliary components function in concert with the core SusCD transporters. Upon addition of FOS, the levan utilisome undergoes large, concerted conformational changes, revealing the substrate-bound states of the Bt1760 levanase and the Bt1761 SGBP, and demonstrating that the pedal-bin mechanism of nutrient import operates in the presence of all OM components of the PUL. Collectively, we show that utilisomes constitute multi-component, macromolecular machines on the cell surface, the architecture of which is consistent with efficient capture and processing of polysaccharides and import of the generated oligosaccharides.

## Results

### Cryo-EM data of the SusCD levan transporter indicate substrate-independent association of auxiliary lipoprotein components

Previously, we used single particle cryo-EM to capture the SusC_2_D_2_ levan transporter produced by fructose-grown *B. theta* in several conformational states (fructose induces expression of the levan PUL)^12^. During 3D classification of these data, compositional heterogeneity of the complex was also identified, with one class (∼10% of the data) possessing density in addition to that expected for the SusCD components (**Fig. 1**). SDS-PAGE and downstream mass spectrometry revealed co-purification of sub-stoichiometric amounts of the levanase (Bt1760) and the levan SGBP (Bt1761). To confirm whether the additional density corresponded to one or both of these components, we docked the crystal structure of *E. coli*-expressed inactive Bt1760 (PDB ID 7ZNR; Supplementary Table 1 and Methods) into the map. The levanase was found to occupy only part of the unassigned density, leading us to tentatively assign the remaining density to Bt1761, for which there was no available structure.

**Figure 1:**
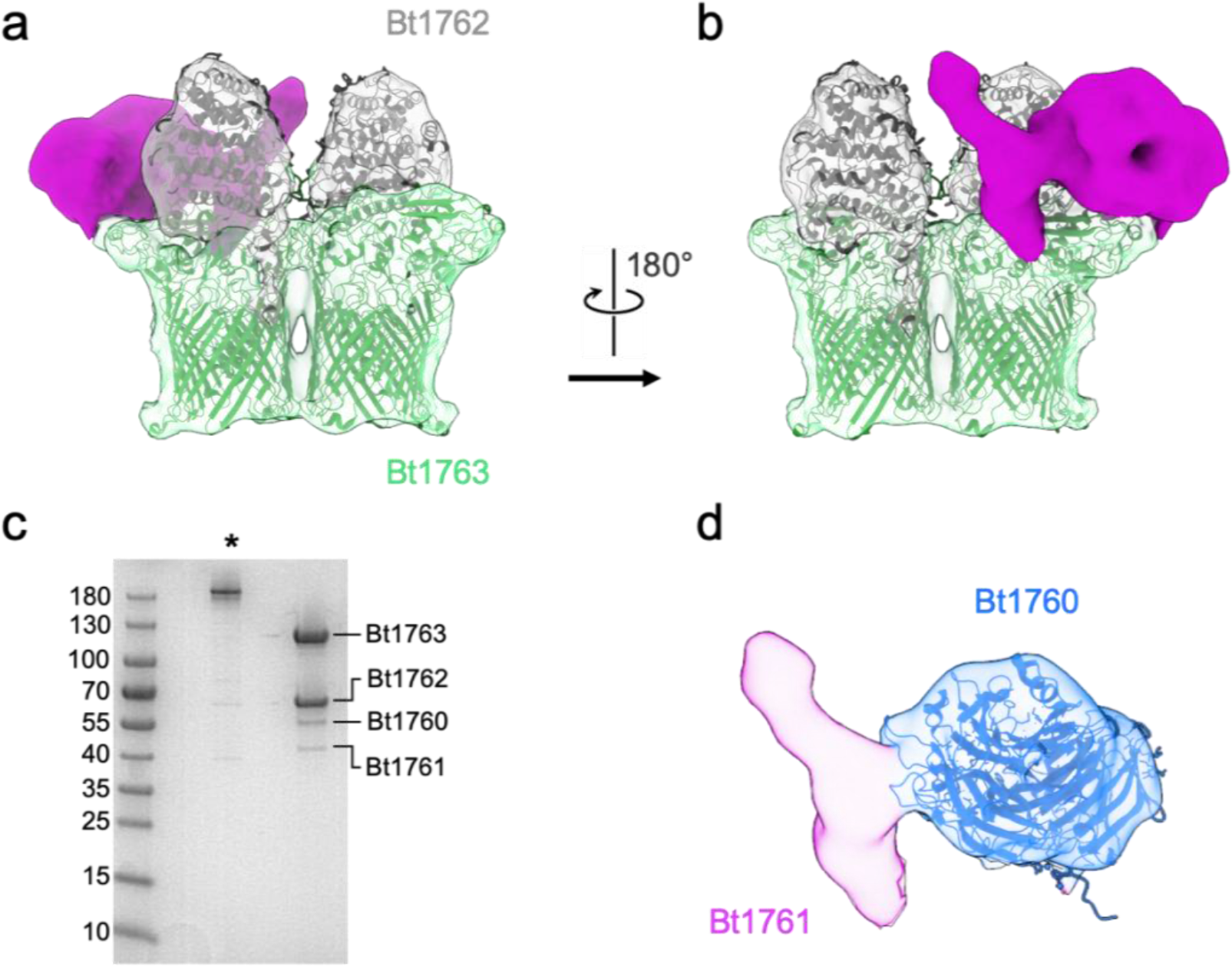
Additional density identified in 3D classification of the levan SusC_2_D_2_ transport complex. **a**, A class average obtained during 3D classification of the levan SusC_2_D_2_ transport complex. The SusC and SusD components (green and grey respectively) are docked into the corresponding density. A large region of density remains unassigned (magenta). **b**, as for **a** rotated 180 degrees. **c**, SDS-PAGE of the sample that was applied to cryo-EM grids before (asterisk) and after boiling. The boiled sample shows two clear bands in addition to those for Bt1762 (SusD) and Bt1763 (SusC), which were identified as Bt1760 levanase and Bt1761 LBP by mass spectrometry. **d**, Isolated view of the previously unassigned density with the crystal structure of Bt1760 (blue cartoon) fit as a rigid body. The structure fits only part of the unassigned density, now coloured blue. The remaining density was therefore attributed to Bt1761 and remains magenta.

The additional density we ascribed to the additional lipoproteins was associated with just one of the two available SusC barrels, and this seemed unlikely to represent a ‘complete’ complex given the dimeric nature of the SusC_2_D_2_ core complex. We therefore adapted the purification protocol to use milder detergents (a mixture of decyl-maltoside, DM and dodecyl-maltoside, DDM) rather than the harsher lauryldimethylamine-oxide (LDAO), to extract the complex from membranes. The remainder of the purification was carried out in DDM rather than in LDAO. In addition, we moved the His_6_-tag from the C-terminus of SusD to the C-terminus of the levan SGBP Bt1761 (Methods). A qualitative assessment of newly purified sample by SDS-PAGE indicated the presence of all four components in approximately stoichiometric amounts (Bt1760-63), within a complex that ran as a single, monodisperse peak in analytical size exclusion chromatography (**Fig. 2a,c**). Similar results were obtained for a dextran-grown *B. theta* strain tagged on the SusD protein from the dextran PUL (Bt3089; **Fig. 2b,c**). In this case, the purified four-component complex (4CC) consisted of Bt3087 (GH66 dextranase), Bt3088 (putative dextran SGBP), Bt3089 (SusD) and Bt3090 (SusC). Structures of the levan and dextran PULs are shown in **ED Figure 1** alongside known and predicted structural information for the OM components.

**Figure 2:**
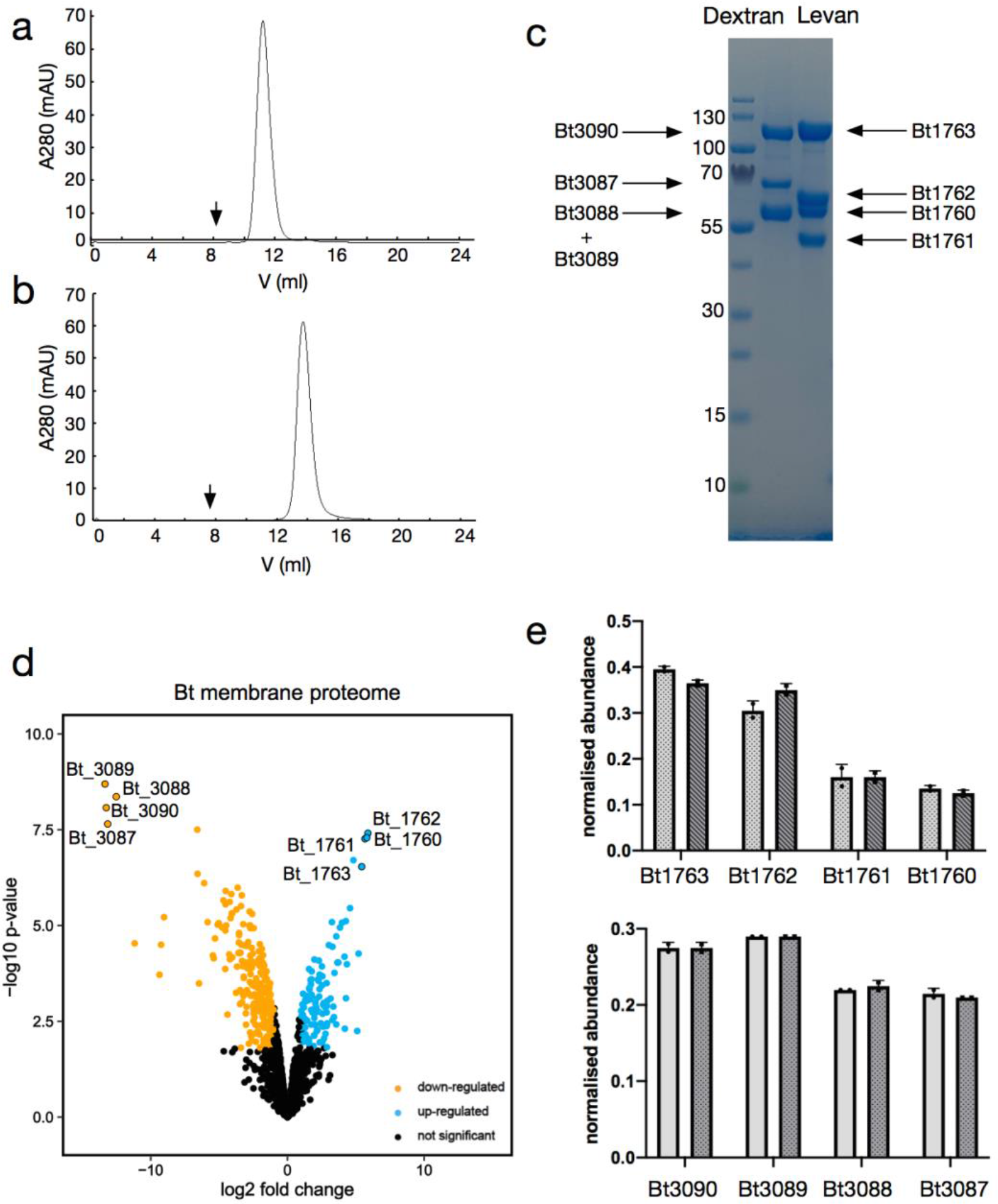
Purification of stable 4-component complexes from the OM of *B. thetaiotaomicron*. **a,b** Analytical SEC elution profiles for Bt1760-63 (levan 4CC; **a**) and Bt3087-90 (dextran 4CC; **b**) analysed on Superdex-200 and Superose-6, respectively. The void volume of the columns are indicated by the arrows. **c**, SDS-PAGE of the complexes purified in (**a,b**) with the identities of the bands indicated. Bt3088 and 3089 of the dextran 4CC are almost the same molecular weight and appear as a single band on the gel. **d**, Volcano plot of the *B. thetaiotaomicron* outer membrane proteome from fructose-vs. dextran-grown cells. **e**, Normalised OM abundance of the four components of the levan (top panel) and the dextran (bottom panel) systems obtained from fructose- and dextran-grown cells, respectively (light bars). The dark bars show the normalised abundance of detergent-purified 4CCs spiked into the proteome samples of dextran- or fructose-grown cells (top and bottom panels, respectively).

### Proteomics data provide support for stable four-component complexes at the OM

Successful purification of two different detergent-solubilised four-component complexes that elute as monodisperse species from size exclusion chromatography strongly suggests that the complexes are stable in the OM. To support this notion and to shed light on the relative abundance of the components, we performed semi-quantitative proteomics using intensity-based absolute quantification (iBAQ)^22^ on the outer membrane fractions from *B. theta* grown on either fructose or dextran. The four components of each complex are among the most-highly expressed membrane proteins in *B. theta* when grown on minimal media with either fructose or dextran as the sole carbon source (**Fig. 2d**). While the proteins of the dextran 4CC are present in roughly equimolar amounts based on the iBAQ analyses, the additional lipoproteins of the levan 4CC appear to be substoichiometric (**Fig. 2e**). However, the SDS-PAGE gel and the monodisperse gel filtration peaks of detergent-purified complex suggest that the iBAQ analysis, being semi-quantitative, may underestimate the abundance of the additional lipoproteins of the levan 4CC.

As expected from transcription studies, the levan PUL is not expressed when *B. theta* is grown on dextran, with the reverse true for expression of the dextran PUL (**Fig. 2d**). Thus, in dextran-grown cells, components of the levan PUL are present at just ∼1 % of the level found when cells are grown on fructose, and in fructose-grown cells, dextran PUL components were not detectable. This provided the opportunity to spike membrane proteome samples from cells grown on fructose or dextran with detergent-purified dextran or levan 4CCs, respectively. For both the levan and the dextran systems, the relative abundance of the four components in the spiked versus endogenous membrane samples was very similar (**Fig. 2e**), suggesting that there is no excess of any component in the OM. Collectively these data support the hypothesis that the additional lipoproteins form stable, equimolar, 4CCs with their cognate SusCD transporters that are dedicated to the generation and import of oligosaccharides, and which we propose to name ‘utilisomes’ in keeping with the ‘PUL’ acronym.

### Single-particle cryo-EM reveals an octameric, four-component complex for levan utilisation

The structure of the DDM-solubilised levan utilisome was assessed by single particle cryo-EM in the absence of substrate. 2D class averages were markedly distinct from those of the core complex, with unambiguous evidence for additional components in the most populated classes (see **ED Figure 2**). Initial classification in 3D revealed extensive compositional heterogeneity in the dataset. Given the biochemical evidence for approximately equimolar complexes, we speculate that this compositional heterogeneity most likely arises from denaturation and dissociation during cryo-EM grid preparation. However, >50 % of the particles represented a complex containing a full complement of components, that is, the SusC_2_D_2_ core, as well as two copies of both the levanase (Bt1760) and the levan SGBP (Bt1761) (see **Figure 3**). We propose that this four-component, octameric complex (∼570 kDa) represents the complete OM levan utilisome. The dataset also contained classes corresponding to partial complexes comprising a single copy of Bt1760 and Bt1761 in addition to the SusC_2_D_2_ core (∼40 % of particles), as well as a small population of ‘naked’ SusC_2_D_2_ complexes (∼12 %).

**Fig 3.**
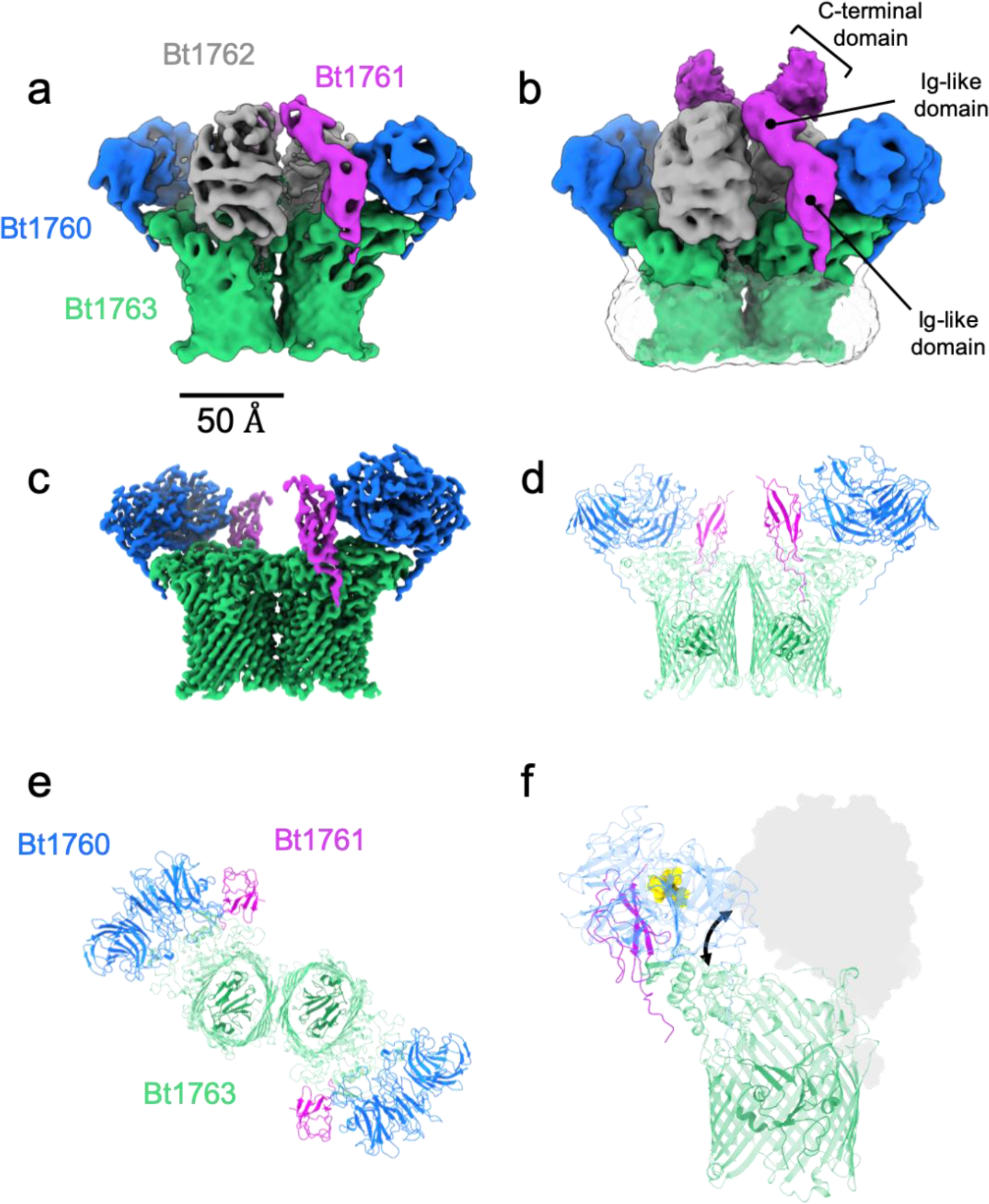
The organisation of the levan utilisome revealed by cryo-EM. **a-b,** Output of 3D refinement for one conformational state of the utilisome obtained by 3D classification. The map is shown at low (**a**) and high (**b**) threshold levels and is viewed in the plane of the membrane. The SusC and SusD components are green and grey respectively. The levan binding protein Bt1761 is shown in magenta and the endo-levanase Bt1760 is coloured blue. Density for the disordered micelle in the high threshold view is transparent grey. **c**, C2 symmetrical reconstruction at 3.5 Å from a focused refinement excluding the variable SusD subunits, viewed in the plane of the membrane. **d**, Atomic model built from the density shown in **c**. **e**, the same model but viewed top down from the extracellular side. **f**, One half of the complex is shown in the context of the SusD lid (grey shadow). Using the crystal structure (PDB ID 7ZNR), FOS DP5 (yellow) has been artificially displayed at the active site of the endo-levanase, highlighting proximity to the open ‘mouth’ of the SusCD transporter (marked by the double headed arrow). Note that all particles possess SusD components, and their absence here is a result of focused refinement.

### The arrangement of the levanase and levan SGBP suggests concerted function

The levanase is mounted on the lip of the SusC barrel, at the opposite edge from the location of the hinge contacts between SusC and SusD (**Figure 3**). Extracellular loops 1 and 9 contribute to the binding interface which buries a surface area of ∼820 Å. This arrangement means that, crucially, the opening and closing of the SusD lid is not hindered by the presence of the levanase. Furthermore, the levanase is oriented in a way that is consistent with concerted function with the core transporter, with its active site facing, and close to, the mouth of the SusCD transporter such that the minimum distance between FOS bound to the respective subunits is ∼30 Å (estimated by alignment of crystal structures of the levanase bound to short FOS (7ZNR) and Bt1763 containing bound FOS (6ZAZ) with the corresponding components in this structure) (**Figure 3**). The levan SGBP is positioned adjacent to the levanase. An AlphaFold2-predicted model^23^ of the levan SGBP has two N-terminal Ig-like domains, followed by a globular C-terminal domain which we postulate is responsible for levan binding, consistent with the domain organisation of other characterised SGBPs (**ED Figure 1**)^24^. In the substrate-free utilisome structure, only the most N-terminal Ig-like domain is unambiguously resolved at high resolution. Interactions between the levan SGBP and SusC again occur at extracellular loop 1. Weaker, poorly-resolved density is present for the second, Ig-like domain, which extends away from the transporter, towards the extracellular space. Finally, the C-terminal domain of the levan SGBP is much more mobile, with weak, diffuse density only visible at high threshold levels (**Figure 3**). The lip of SusC therefore serves as a platform for the association of the auxiliary components, perhaps explaining why SusC-type TonB-dependent transporters are ∼40 % larger than their classical monomeric counterparts^9^. The structure, alongside the observation that most of the levan SGBP subunits appear to be flexible, provides important insight into how the utilisome architecture contributes to efficient levan utilisation. The putative levan binding C-terminal domain of the levan SGBP is projected away from the cell surface, facilitating initial capture of levan chains. The observed flexibility of this region would then feasibly permit positioning of these chains for processing by the adjacent levanase units. Finally, any cleavage products would be released close to their binding site at the SusCD interface, promoting efficient loading of the transporter.

### Conformational heterogeneity of the apo utilisome complex

We next performed further 3D classification of particles corresponding to the intact utilisome. The major area of conformational heterogeneity within these data is in the position of the SusD lids (**ED Figure 3**). All complexes within this dataset are open, but they are open to differing degrees, suggesting a rugged energy landscape for the lid-open state with several local minima. The lids within a complex appear to operate independently of one another such that differing lid positions on either side of the complex break its C2 symmetry. The lid conformations vary from a partially open state that is comparable to the predominant lid position observed in the study of the core apo SusC_2_D_2_ complex^12^, to a wide-open state also reported previously (**ED Figure 3**). Importantly, no closed lids are observed and, in contrast to our findings for the LDAO-solubilised SusC_2_D_2_ core complex^10, 12^, all SusC barrels contain unambiguous density for the SusC plug domain that occludes the barrel. Thus, transporters with open lids are plugged and competent to accept substrate, consistent with the proposed ‘pedal-bin’ hypothesis, Mobility of the C-terminal region of the levan SGBP is also evident in the output of 3D classification (**ED Figure 3**).

Whilst the SusD lids have several conformations, SusC, the levanase and the N-terminal domain of the levan SGBP constituted a rigid unit with C2 symmetry. To provide more detailed structural information for these regions, a mask that excluded density for the SusD lids was applied in a focused refinement, generating a reconstruction at 3.5 Å after sharpening (**Figure 3**). At this resolution, it was possible to dock and refine models for SusC and the levanase, and to build the N-terminal domain of the levan SGBP. The refined model of Bt1760 within the utilisome is similar to the crystal structure of *E. coli*-expressed Bt1760 solved in isolation (PDB: 7ZNR)(Cα rmsd = 0.54 Å).

### A stable, four-component utilisome is not unique to the levan utilisation system

To investigate whether the levan utilisome architecture is a general feature of OM PUL components, we studied a second example; the dextran utilisation system of *B. theta*. Dextran is an α1,6 linked glucose polymer with occasional α1,4 glucose branches and, like the levan utilisation system, the dextran PUL encodes 4 OM components: the SusC transporter and SusD lid (Bt3090 and Bt3089), a GH66 endo-dextranase (Bt3087), and a putative dextran SBGP (Bt3088) (**ED Figure 1**).

Cryo-EM of the dextran system revealed it to be more aggregation prone than the levan system, requiring altered grid preparation conditions for data collection (Methods). Despite this, initial 3D classification shows that the SusC_2_D_2_ complex is intact with SusD subunits occupying an open position. As observed for the levan system, there is considerable compositional heterogeneity in the data. Densities for the dextranase and dextran SGBP were first assigned by rigid-body docking of AlphaFold2-predicted models (**ED Figure 4).** Both lipoproteins were observed to assemble on the core SusC_2_D_2_ transporter although, unlike the levan system, there were no classes corresponding to an octameric dextran utilisome, which we attribute to a higher propensity of components of the dextran system to disassemble/aggregate during grid preparation. The highest order complex observed was heptameric, containing two copies of the dextran SGBP and a single dextranase. Hexameric and pentameric complexes comprised of all possible complements of additional lipoproteins (**ED Figure 5**) were also observed. Interestingly, the relative positions of the auxiliary components on the core complex differ between the levan and dextran utilisomes (**ED Figure 6**). For the former, the endo-levanase is nearest to the hinge of SusD whereas for the latter the glycan binding protein is nearest. Regardless, the arrangement of components in the dextran utilisome is compatible with movement of the SusD lid. Together, the data suggest that OM utilisomes are a general feature of *Bacteroides* PULs.

### Addition of levan causes concerted conformational changes in the utilisome

To probe the mechanism of substrate capture and processing by the levan utilisome, ∼0.5 mM levan FOS with a degree of polymerisation (DP) of ∼8-12 (DP8-12) was added to the detergent-purified utilisome complex. The resulting 3D structure shows that a major, concerted conformational change has occurred in the complex in the presence of FOS, with all SusD lids closed and tightly capping the lumen on the extracellular side of the SusC barrels (**Figure 4**). 3D classification of these data suggested that all major heterogeneity was due to the presence/absence of auxiliary lipoproteins (**ED Figure 7**). To confirm the presence or absence of substrate within the closed transporter complex, we first performed targeted refinement of the SusC_2_D_2_ core. To achieve this, we used particle subtraction to remove the signal for the levanase and levan SGBP from the images, and were thus able to include all particles in our refinement, regardless of their lipoprotein complement. A final reconstruction of the SusC_2_D_2_ core was obtained at 2.9 Å, and contains unambiguous density for FOS substrate at two distinct sites within the SusC barrel. These densities are comparable to those observed in the crystal structure of the substrate-bound SusC_2_D_2_ complex (6ZAZ) (**Figure 4**). The oligosaccharide bound at the interface of SusC and SusD consists of six β2,6-linked fructose units with a β2,1 decoration on Frc-4 (numbered from the non-reducing end). The FOS adopts a compact, twisted topology similar to that observed for the FOS in the SusC_2_D_2_ crystal structure (6ZAZ) and FOS DP5 in the levanase crystal structure. The second FOS molecule occupies a site at the bottom of the SusC binding cavity, where it is in contact with the plug. Here the density most likely corresponds to four β2,6-linked fructose units.

**Fig 4.**
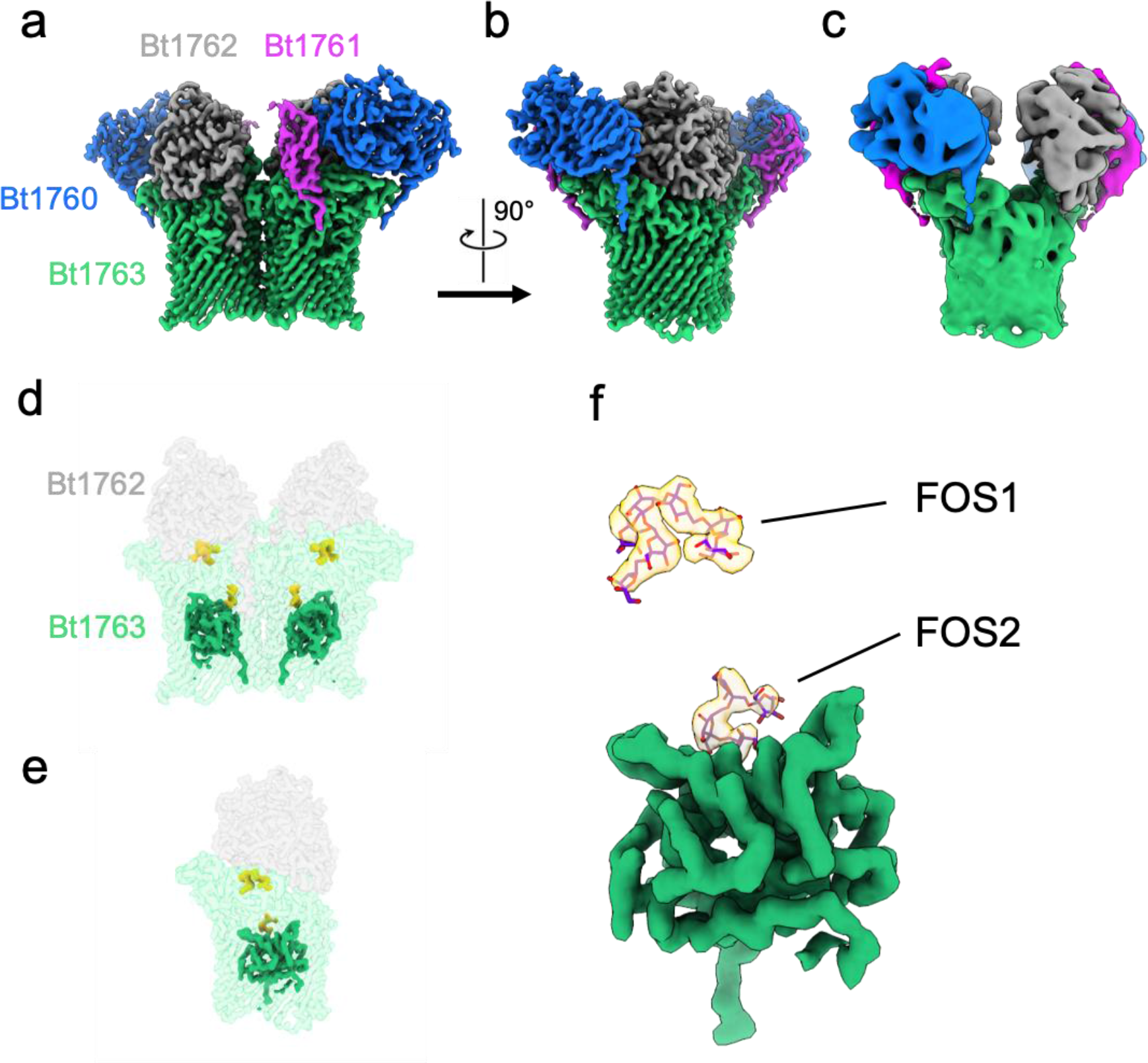
Cryo-EM shows a concerted conformational change of the levan utilisome upon addition of short FOS. **a-b**, Reconstruction at 3.2 Å of the closed substrate-bound transporter viewed in the plane of the membrane **(a)** and rotated 90° (**b**). As before, SusC is coloured green, SusD grey, levan SGBP magenta and levanase blue. **c**, Wide open state of the dimeric transporter seen in the absence of substrate in same orientation as in b, shown for comparison. **d**, Reconstruction of the SusC_2_D_2_ core complex generated from a focused C2 refinement after particle subtraction of additional lipoproteins. Densities for bound FOS (yellow) and the plug domain (solid green) of SusC are highlighted. **e**, Isolated view of a single SusCD subunit from (**d**) shown from the dimerisation interface. **f,** Isolated view of plug and substrate density in the same orientation as (**e**). The FOS modelled are similar to those in the previously determined crystal structure of SusC_2_D_2_ (PDB: 6ZAZ). Maps for the substrate-bound structure are LAFTER filtered (see **Methods**).

Given the biological role of the utilisome, it is expected that substrate loading of the transporter should be signalled across the OM. This would presumably occur via disordering of the SusC Ton box region, at the periplasmic face of the complex^12^. Indeed, the Ton box and the region preceding it are not visible in the substrate-bound structure (**ED Figure 8**). This is in stark contrast to the apo state of the transporter in which the region preceding the Ton box is resolved, and adopts a sharp turn towards the plug domain, constraining the Ton box within the barrel. It should be noted that the N-terminal extension (NTE) domain of SusC, that we have previously shown to be essential for function^12^, is not resolved in either the apo or substrate-bound states of the complex.

The positions of Bt1763 substrate binding residues are almost identical in the apo and FOS-bound cryo-EM structures (**ED Figure 9**). Therefore, the conformational changes upon substrate binding that lead to exposure of the Ton box are not obvious. One notable difference between the apo and substrate-bound structures is the position of Hinge 2 (or L8) of SusC^12^. Hinge 2 forms extensive contacts with SusD and interacts with FOS at the top of the SusC cavity via F649 (**ED Figure 9a**). In our apo structure, most of Hinge 2 is not visible, likely because it is masked out together with the flexible SusD lid during data processing (Methods). However, the part of Hinge 2 proximal to the SusC barrel is resolved in both structures and shows differences in the side chain orientations which propagated along the edge of the barrel and plug towards the periplasm (**ED Figure 9**). In the substrate-bound structure, W685, which sits at the base of Hinge 2, shifts down towards the periplasm, nudging F583 up and S193 down. This results in a downward shift of the plug loop connecting α3 and β6, to which S193 belongs. In the apo structure, Y191, which is also part of the α3-β6 plug loop, forms a triple aromatic stack together with Y89 and F588. When the α3-β6 plug loop shifts towards the periplasm, the stacking interaction between Y191 and Y89 is broken, and Y191 clashes with the C-terminus of the Ton box resulting in its release into the periplasm. Recent studies of *E. coli* BtuB (vitamin B TBDT) highlighted the presence of a salt bridge between the transporter barrel and plug residues, which has been termed an ‘ionic lock’^25^, that is broken upon vitamin B12 binding^26^. Similarly, we propose to name the Bt1763 Y89-Y191 stacking interaction an ‘aromatic lock.’ Our cryo-EM structures reveal how conformational changes caused by substrate-induced SusD lid closure on the extracellular side of the outer membrane propagate through the TonB-dependent transporter, breaking the aromatic lock on the periplasmic side and exposing the Ton box.

As expected, no substrate density was observed within the active site of the wild type levanase. The addition of short FOS substrate had no influence on the conformations of the auxiliary lipoproteins, and the flexibility of the levan SGBP prevented further structural analysis. Notably, in both the apo and substrate-bound datasets, the levanase and levan SGBP were invariably associated with one another, i.e. there is no evidence of pentameric or heptameric complexes where one SusCD unit is associated with a single lipoprotein as is the case for the dextran system.

### Longer substrate chains tether the levan SGBP

While the cooperative behaviour of the levan utilisome components is implied by their arrangement, the structure of the complete levan binding protein remained unresolved. We reasoned that by using a catalytically inactive levanase (with the active site D42A mutation) and longer FOS chains (DP ∼15-25), cooperative binding of a single levan chain by the levan SGBP and the levanase might restrict the mobility of the levan SGBP. We therefore collected a new cryo-EM dataset for the utilisome with inactivated levanase and long-chain FOS, and observed that in 3D classifications, a novel conformation of the levan SGBP was present. In several classes, the levan SGBP adopted a ‘docked’ state, in which its C-terminal domain was held proximal to both SusD and the levanase (**ED Figure 10**). However, a consensus refinement of those classes containing evidence of the docked state still produced a map in which the C-terminal domain of Bt1761 was poorly resolved (**ED Figure 10**), indicating that local conformational heterogeneity remained. We therefore attempted to collate a more homogeneous subset of particles from within the dataset by using an approach involving focused classification without particle alignment (Methods), using a mask that only included the docked levan SGBP position. This classification yielded a class with high-resolution information for the C-terminal domain of the levan SGBP (**ED Figure 10**). Particles belonging to this class were used to generate a final, sharpened reconstruction of the complete utilisome at 3.0 Å with well-resolved density for the complete levan SGBP (**Figure 5**), which we built into the cryo-EM map, starting from a partial model obtained by X-ray crystallography (Methods).

**Fig 5.**
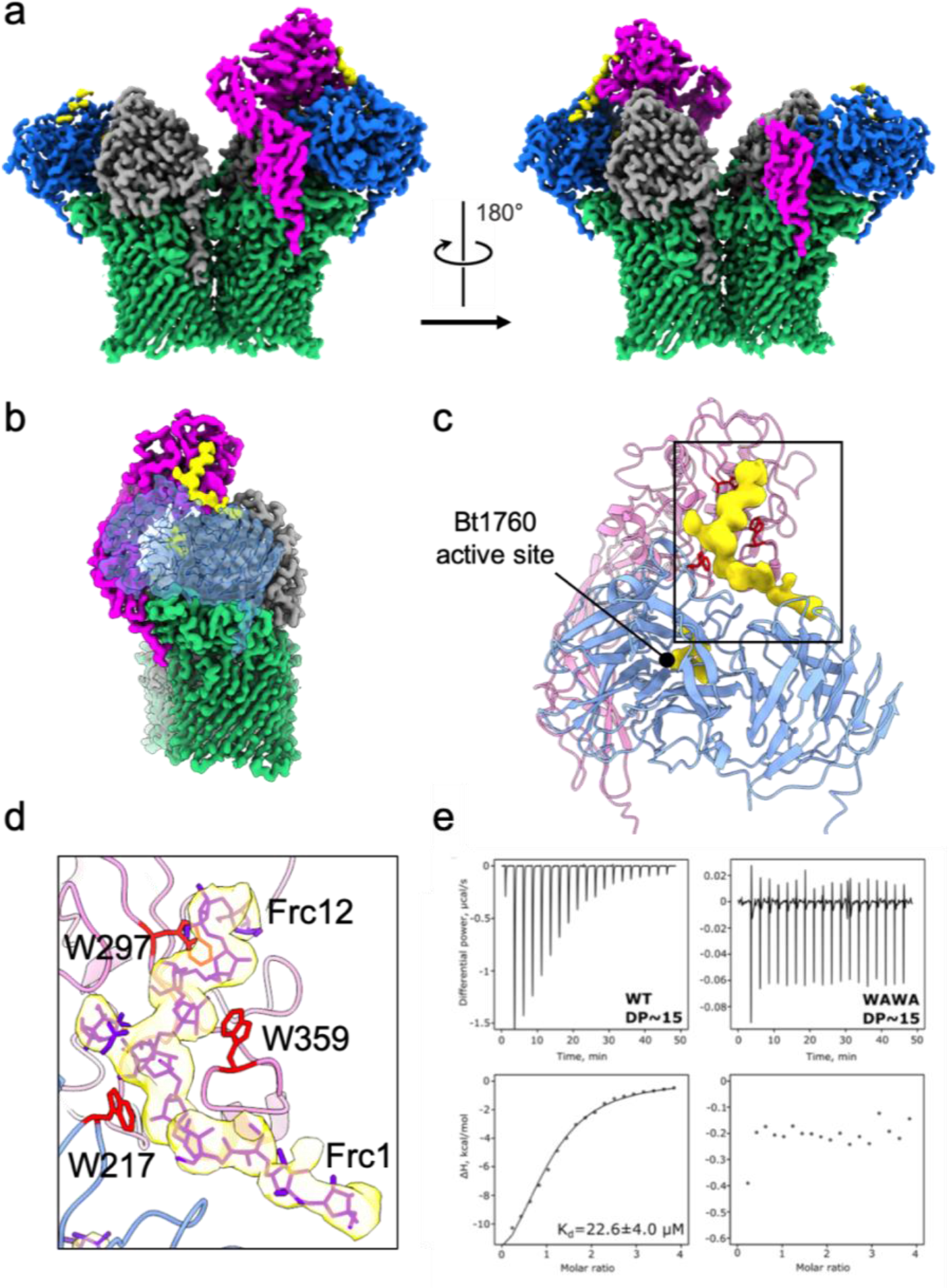
Cryo-EM structure of the holo utilisome. **a**, Reconstruction of the levan utilisome complex after focused classification viewed in the plane of the membrane (left) and rotated 180° (right). One levan SGBP is now fully resolved (magenta). **b**, side-on view of the complex with transparent density for Bt1760, highlighting the position of the cooperatively bound levan chain (yellow). **c**, Structures of the levan SGBP and inactive endo-levanase with additional density attributed to the cooperatively bound FOS shown in yellow (boxed). **d**, Zoomed inset of the boxed region in **c**. The modelled levan chain is shown as purple sticks. Tryptophan residues proposed to contribute to binding are shown in red and from top to bottom correspond to W297 and W359 from Bt1761 levan SGBP and W217 from Bt1760 levanase. Note that the FOS that bridges the levan SGBP and the levanase occupies a secondary binding site on the latter, distinct from the active site. Maps displayed here have been filtered using LAFTER^27^. **e**, Isothermal titration calorimetry experiments with 1 mM FOS DP∼15 titrated into 50 μM recombinant wild type levan SGBP, or the W297A/W359A double-mutant (WAWA). Removing both tryptophan residues completely abolished binding.

In agreement with the AlphaFold2 structure prediction, the levan SGBP is organised into three domains, with the lipidated N-terminal and central domain each possessing an Ig-like fold. The C-terminal levan-binding domain (residues 205-321) is globular, with a central β-sheet decorated with α-helices. A distance-matrix alignment (DALI) analysis of the C-terminal domain returned a number of structures with significant similarity. A ThuA-like protein (PDB ID 1T0B) with a putative role in the conversion of disaccharides to their 3-keto derivatives and a homoserine O-succinyl transferase (PDB ID 7CBE), have Z-scores of ∼15 and ∼14 respectively.

Additional density forms a bridge between the levanase and levan SGBP. We attribute this density to a length of levan chain, bound by the C-terminal domain of the levan SGBP and the inactive levanase. The density is compatible with a stretch of 12 β2,6-linked fructose units, with a putative β2,1 decoration observed on Frc-7 (numbered from non-reducing end) (**Figure 5**). Interestingly, the bound levan chain adopts a relatively extended conformation, markedly different from the compact twisted conformations observed for FOS bound to the levanase and within the SusCD units. Whilst the resolution of the density is insufficient for a detailed description of binding interactions, several tryptophan residues are clearly implicated in binding. These include W297 and W359 from the levan SGBP, and W217 from the levanase, which appear to cradle the FOS chain via stacking interactions. Isothermal titration calorimetry (ITC) on recombinant protein produced in *E. coli* confirmed the importance of W297 and W359 for levan binding by the levan SGBP (**Figure 5e, ED Figure 11**). In addition, the ITC data suggest that the binding site observed in the cryo-EM structure is the only levan binding site on the levan SGBP. The minimum FOS size that the levan SGBP can bind reasonably strongly is an 8-mer (**ED figure 11)**.

Interaction of the long, bridging, levan chain with the levanase does not occur at the levanase active site. Instead, the bridging levan binds across the top of the β-propeller domain of the enzyme and the levan density then proceeds towards the C-terminal β-sandwich domain (**Fig. 5, ED Figures 12, 13**). Evidence of levan binding at this secondary ‘tethering’ site on the levanase is also seen in the absence of bridging interactions with the levan SGBP and in crystal structures of the inactive levanase (D42N) solved in the presence of FOS of DP ∼7-8 (**ED Figure 13)**. Indeed, the presence of a secondary binding site (SBS) is relatively common in endo-acting GHs and is proposed to be involved in enhancing enzyme-substrate targeting and processivity. Notably, no ligand was observed at the secondary site in a levanase structure complexed to levantetraose (PDB ID 6R3U; ^28^), suggesting the affinity of this site for very short FOS is low. The mutated active site of the enzyme is occupied by a distinct FOS molecule, with density for five β2,6 linked fructose units that adopt the same twisted conformation as levantetraose in 6R3U (**ED figure 13**). No obvious levan density is observed between the tethering site and active site.

We attempted to dissect the affinities of the levanase active site and the SBS for levan. The protein-FOS interactions are extensive in both binding sites and include numerous aromatic residues that are often found at carbohydrate interaction interfaces (**ED Figure 14**). We tested the effects of substituting these aromatic residues to alanine on levan binding (**ED Figure 15**). Substitutions near the active site (Y70A and W318A) decreased the affinity for levan 25-30 fold. Single substitutions in the SBS either had no effect (F243A and Y437A) or reduced the affinity 6-fold (W217A). These results suggest the residues near the active site are responsible for most of the affinity of the levanase for levan, and the SBS provides a modest contribution. However, it is possible that the numerous weak interactions of the SBS with levan, including polar residues, result in a substantial overall affinity.

Why SusCD transporters exist as dimers is one of the key unanswered questions in the field. While the functional relevance of dimerisation is not clear when thinking about the core SusC_2_D_2_ complex in isolation, considering the question in the context of the utilisome provides some important insights. During 3D classification of both the active and inactive substrate-bound utilisome datasets, the inherent flexibility of the levan SGBP was evident. However, in the inactive utilisome dataset with long FOS, a novel conformation was observed in which an untethered levan SGBP appears to reach across and contact a tethered levan SGBP associated with the other SusC subunit (**ED Figure 16**). The fact this ‘reach over’ conformation was observed exclusively in the inactive utilisome dataset with the longer FOS perhaps indicates that this conformation is the result of both levan SGBPs interacting with the same stretch of the long levan chain that would likely be present *in vivo*. Such cooperative binding of long glycan chains by the levan SGBPs would result in higher avidity for the substrate and would bring/keep more substrate in the vicinity of the utilisome, offering a potential explanation as to how dimerisation contributes to function.

### FOS binding by the SusCD core is promiscuous

Interestingly, in the presence of longer FOS and an inactive levanase, the substrate density within the SusCD binding cavity differs from that observed previously for short FOS. At the FOS1 site, density for the putative β2,1 decoration on Frc4 is missing (**ED Figure 17)**. Conversely, contiguous density can be seen beyond the previously defined reducing terminus of FOS1, with a novel β2,1 decoration on Frc5. The substrate bound at the FOS2 site follows a similar trend with the β2,1 linked fructose side chain modelled previously being much weaker with longer FOS, and additional density attributed to another β2,6 linked monomer extending the chain in the direction of the FOS1 site. Strikingly, at higher threshold levels, diffuse density is visible that connects both FOS binding sites, indicating that longer FOS (∼DP15) can occupy both sites simultaneously. The density between the FOS1 and FOS2 sites is relatively weak and indicative of multiple conformations, consistent with the absence of any contacts from SusC in this bridging segment. These data confirm that the transporter has considerable substrate promiscuity and that, as suggested previously, relatively long FOS (∼15 DP) can be accommodated^12^.

## Discussion

Our cryo-EM structures of the levan utilisome provide unprecedented insight into glycan acquisition by *Bacteroides spp*. Since all sequenced Bacteroidetes to date contain SusCD homologues, our findings likely apply across multiple systems, potentially including SusCD systems involved in the transport of non-glycan macromolecules^10^. As Bacteroidetes are found in a diverse range of terrestrial and marine niches^29^, the data also expand our understanding of glycan utilisation outside the animal gut. We demonstrate that for two distinct PULs, all of the gene products that localise to the bacterial OM form a stable utilisome complex with a defined architecture. The position of each component within this complex is consistent with their respective functions, encapsulated in our general model for utilisome function (**Fig. 6**). In the absence of substrate (i), the SGBP is in an extended conformation with a mobile glycan-binding domain, increasing its efficiency as a glycan grappling device. Likewise, the SusD lids, which can open and close without clashing with the other components, are mobile and occupy predominantly open states. (ii) Glycan binding by the SGBP is followed by docking to the proximal glycanase and binding by the SBS and active site of the enzyme. Substrate cleavage by the endo-acting enzyme generates shorter oligosaccharides in close vicinity to the mouth of the SusCD transporter. (iii) Concerted binding of the oligosaccharide to SusD and SusC promotes the formation of the closed conformation of the transporter, leading to conformational changes along the SusC barrel and plug, breaking of the aromatic lock and exposure of the Ton box to the periplasm. This is followed by TonB-dependent transport events (not shown), and resetting of the transporter to the open state (iv)^10, 12^.

**Figure 6.**
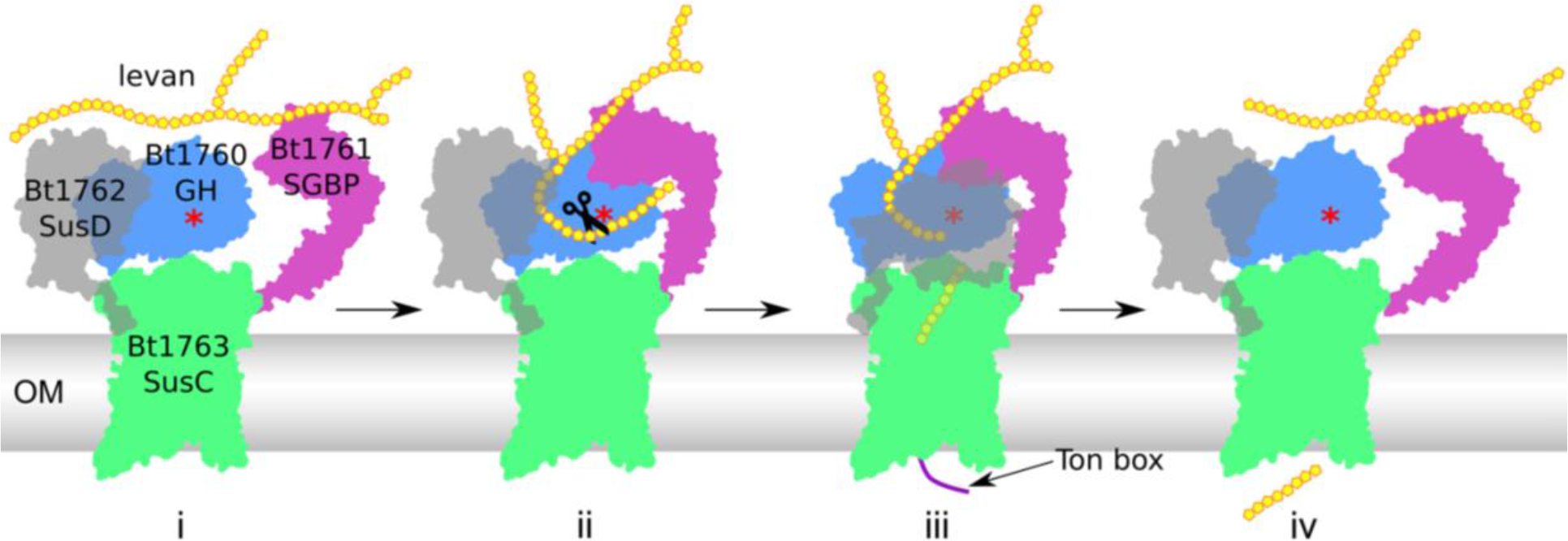
The transport cycle of glycan utilisomes. Description of states i-iv is provided in the main text. N.B. only one half of the dimeric utilisome is shown for clarity. The red asterisk marks the active site of the glycoside hydrolase (GH). OM, outer membrane.

The bridge formed by long levan chains (DP ∼15-25) between the levan binding domain of the SGBP and the levanase raises several interesting points. As outlined above, its extended conformation is unique, perhaps arising because the chain is held under tension. The utilisome structure captured here may therefore represent a state that is ‘strained’, with a bound levan chain that is pulled taut. However, it is important to note that this conformation has been selected for during image processing owing to the fact that it represents the least mobile and therefore most resolvable state. Second, even though the SBS of the levanase is ∼25 Å away from its catalytic site, we speculate that the levan SGBP-levanase interaction serves to increase the local levan concentration around the active site, optimising the efficiency of substrate cleavage. Flexibility of the levan chain away from the interaction sites may preclude resolution of contiguous density in our structure or alternatively, the FOS present in our sample are not long enough to bridge both sites.

Whilst our data argue in favour of stable utilisome assemblies in the OM, recent studies on the archetypal starch PUL propose a much more dynamic model, with immobile starch binding centres comprised of the SGBPs SusE and SusF, around which transporter components and the endo-amylase (SusG) can transiently assemble^21^. Proteomics data for the starch utilisation system reveal similar amounts of SusC and SusD at the OM whilst SusEFG are much less abundant, providing support for the presence of complexes with differing stoichiometries^21^. Intriguingly, co-immunoprecipitation with SusD antibodies captures twice as much SusD as SusC, suggesting that even the core SusCD transporter may not form a stable complex in the starch system. This clearly contrasts with our work, where separate SusC or SusD components have never been observed during purification, regardless of whether the purification tag was located on SusC or SusD. Moreover, while it is not clear how much the *E. coli* OM resembles that of *B. theta*, recent work has shown that *E. coli* OMP and LPS mobility is limited ^30, 31^, raising questions regarding the efficiency of dynamically (dis)assembling transport complexes in a crowded OM.

It is not clear why the starch system appears to operate differently to the levan and dextran systems presented here. One consideration is that these models are based on live-cell fluorescence studies with C-terminal fusions of OM PUL components with relatively large fluorescent protein tags. Considering the compact utilisome architecture (**ED Figure 4**), it is possible that the fluorescent tags cause utilisome destabilisation or interfere with assembly, resulting in dynamic behaviour that might not affect cell growth *in vitro*. On the other hand, the starch PUL is different as it encodes two SGBPs (SusE and SusF), *i.e*. there are five OM PUL components and not four as in the systems studied here. An AlphaFold2 structure prediction of the starch SusC (Bt3702) shows that it has no extra interaction surface for the association of additional lipoprotein components relative to the levan and dextran SusC units (**ED Figure 6**), making it unlikely that SusE, SusF and SusG can all bind the same SusCD transporter and suggesting the starch utilisome may be more dynamic.

A key remaining question is how utilisomes are assembled. Surface lipoproteins, including the SusD-like components, glycoside hydrolases and SGBPs, must be flipped from the inner to the outer leaflet of the OM for presentation on the cell surface. The size and complexity of these subunits suggests that dedicated cellular machinery is required, although candidates have yet to be identified. It is conceivable that this process is coordinated with the folding and insertion of SusC components into the OM, in which case the β-barrel assembly machinery (BAM) would be closely associated with the putative lipoprotein flippase.

## Author contributions

JBRW performed cryo-EM and determined cryo-EM structures, supervised by NR. AS purified proteins, determined structures and carried out ITC. MF purified proteins. YL prepared OM samples for proteomics. TH and AG performed proteomics, supervised by MT. HZ purified Bt1760, supervised by DNB. SF collected X-ray crystallography data for Bt1760. AB solved Bt1760 crystal structures and managed the Newcastle Structural Biology Laboratory. BvdB generated *B. theta* strains, purified proteins, and crystallised Bt1761. BvdB coordinated the project. JBRW, AS, DNB, NAR and BvdB wrote the manuscript.

## Data availability

The data supporting the findings of this study are available from the corresponding authors upon reasonable request. Cryo-EM reconstructions and corresponding coordinates have been deposited in the Electron Microscopy Data Bank and the Protein Data Bank respectively: Substrate free levan utilisome (EMD-15288, PDB ID 8A9Y), levan utilisome with FOS DP 8-12 (EMD-15289, PDB ID 8AA0), SusC_2_D_2_ core from levan utilisome with FOS DP 8-12 (EMD-15290, PDB ID 8AA1), Inactive levan utilisome with FOS DP 15-25 (EMD-15291, PDB ID 8AA2), SusC_2_D_2_ core from inactive levan utilisome with FOS DP 15-25 (EMD-1592, PDB ID 8AA3), dextran utilisome consensus refinement (EMD-15293, PDB ID 8AA4). Raw cryo-EM movies will be deposited in the Electron Microscopy Public Image Archive. Coordinates and structure factors from X-ray crystallography experiments that support the findings of this study have been deposited in the Protein Data Bank under the accession codes 7ZNR and 7ZNS. The mass spectrometry proteomics data have been deposited to the ProteomeXchange Consortium via the PRIDE partner repository with the dataset identifier PXD034863.

## Acknowledgements

J.B.R.W. was supported by a Wellcome Trust 4-year Ph.D. studentship (215064/Z/18/Z). B.v.d.B is funded by a Wellcome Trust Investigator award (214222/Z/18/Z), supporting AS and YL. MF is supported by a Newcastle University studentship. We acknowledge the Diamond Light Source (Didcot, UK) for beam time (proposals mx306, mx1221, mx13587 and mx18598), and thank the staff of beamlines I02, I03, I04 and I24 for support. All cryoEM was performed at the Astbury Biostructure Laboratory, which was funded by the University of Leeds and Wellcome (108466/Z/15/Z, 221524/Z/20/Z). We thank R. Thompson, E. Hesketh and D. Maskell for EM support. Protein ID mass spectrometry was performed at the University of Leeds Mass Spectrometry Facility. We thank J. Ault and R. George for performing this analysis. For the purpose of open access, the authors have applied a CC BY public copyright licence to any Author Accepted Manuscript version arising from this submission.

## Methods

### Construction of *B. theta* strains

*B. theta* strains were made as described previously ^12^. Briefly, the DNA sequence containing the desired alterations was constructed using the sewing PCR method and ligated into the pExchange-tdk vector^32^. The vectors carrying the altered DNA sequences were introduced into the *B. theta tdk^-^*strain via conjugation from *E. coli* S17 λ *pir*. Chromosomal alterations were made by allelic exchange, followed by selection for loss of the pExchange-tdk vector backbone. Mutations were confirmed by PCR amplification of the region of interest and Sanger sequencing.

### Expression and purification of utilisomes from *B. theta*

*B. theta* strains were grown at 37°C in a Don Whitley Scientific A32 anaerobic workstation. Brain-heart infusion (BHI) cultures supplemented with 2 ug/ml hemin were inoculated with stabs from glycerol stocks of the appropriate *B. theta* strain and incubated overnight. Defined minimal medium^12^ was supplemented with 2 ug/ml hemin and either 0.5% fructose (levan system) or 0.5% dextran (dextran system) and inoculated with the overnight BHI cultures (1:1000 dilution). The cultures were harvested after 20 h by centrifugation and the pellets were stored at −20°C. 4 litres of culture were grown for Bt1762-His (with wild type levanase) and Bt1761-His/Bt1760-D42A (inactive levanase) strains for cryo-EM.

The cell pellets were thawed, supplemented with DNase I and homogenised in Tris-buffered saline (TBS, 20 mM Tris-HCl pH 8.0, 300 mM NaCl). The cells were lysed with a single pass at 22 kpsi through a cell disruptor (0.75 kW; Constant Systems). Membranes were isolated by ultracentrifugation for 45 min at 42,000 rpm (45 Ti rotor, Beckman), 4°C. The membranes were solubilised at 4°C for 1 h in TBS with 1% DDM while stirring. Insoluble material was pelleted by ultracentrifugation for 30 min at 42,000 rpm (45 Ti rotor), 4°C. The supernatants were supplemented with 20 mM imidazole and loaded onto an 8 ml chelating sepharose column charged with Ni^2+^, by gravity flow at room temperature. The column was washed with 20 column volumes TBS with 30 mM imidazole and 0.15% DDM. The bound proteins were eluted with 3 column volumes TBS with 250 mM imidazole and 0.15% DDM. The eluates were concentrated in an Amicon Ultra filtration device with a 100 kDa cut-off membrane. The samples were then loaded on a HiLoad 16/600 Superdex 200 pg column (Cytiva) in 10 mM HEPES-NaOH pH 7.5, 100 mM NaCl, 0.03% DDM. Fractions containing pure protein were pooled, concentrated, flash-frozen in liquid nitrogen and stored at −80°C.

### Sample preparation for mass spectrometry

*Bacteroides thetaiotaomicron* membrane samples were suspended in 5% sodium dodecyl sulfate (SDS) in 50 mM triethylammonium bicarbonate (TEAB) pH 7.5 and sonicated using an ultrasonic homogenizer (Hielscher) for 1 minute. Samples were centrifuged at 10,000 xg for 10 minutes to pellet debris. Proteins (40 µg) were subsequently reduced by incubation with 20 mM tris(2-carboxyethyl)phosphine for 15 minutes at 47 °C, and alkylated with 20 mM iodoacetamide for 15 minutes at room temperature in the dark. Proteomic sample preparation was performed using the suspension trapping (S-Trap) sample preparation method^33, 34^, with minor modifications as recommended by the supplier (ProtiFi™, Huntington NY). Briefly, 2.5 µl of 12% phosphoric acid was added to each sample, followed by the addition of 165 µl S-Trap binding buffer (90% methanol in 100 mM TEAB pH 7.1). The acidified samples were added, separately, to S-Trap micro-spin columns and centrifuged at 4,000 xg for 1 minute until all the solution has passed through the filter. Each S-Trap micro-spin column was washed with 150 µl S-trap binding buffer by centrifugation at 4,000 xg for 1 minute. This process was repeated for a total of five washes.

Twenty-five µl of 50 mM TEAB containing 4 µg trypsin was added to each sample, followed by proteolytic digestion for 2 hours at 47 °C using a thermomixer (Eppendorf). Peptides were eluted with 50 mM TEAB pH 8.0 and centrifugation at 3,000 xg for 1 minute. Elution steps were repeated using 0.2% formic acid and 0.2% formic acid in 50% acetonitrile, respectively. The three eluates from each sample were combined and dried using a speed-vac before storage at −80°C.

### Mass spectrometry

Peptides were dissolved in 2% acetonitrile containing 0.1% formic acid, and each sample was independently analysed on an Orbitrap Q Exactive HF mass spectrometer (Thermo Fisher Scientific), connected to an UltiMate 3000 RSLCnano System (Thermo Fisher Scientific). Peptides were injected on a PepMap 100 C18 LC trap column (300 μm ID × 5 mm, 5 μm, 100 Å) followed by separation on an EASY-Spray nanoLC C18 column (75 μm ID × 50 cm, 2 μm, 100 Å) at a flow rate of 250 nl/min. Solvent A was water containing 0.1% formic acid, and solvent B was 80% acetonitrile containing 0.1% formic acid. The gradient used for analysis was as follows: solvent B was maintained at 2% B for 5 min, followed by an increase from 2 to 30% B in 110 min, 30% to 42% B in 10 min, 42-90% B in 0.5 min, maintained at 90% B for 4 min, followed by a decrease to 2% in 0.5 min, and equilibration at 2% for 20 min. The Orbitrap Q Exactive HF was operated in positive-ion data-dependent mode. The precursor ion scan (full scan) was performed in the Orbitrap (OT) in the range of 350-1,500 m/z with a resolution of 60,000 at 200 m/z, an automatic gain control (AGC) target of 3 × 10^6^, and an ion injection time of 50 ms. MS/MS spectra were acquired in the OT using the Top 20 precursors fragmented by high-energy collisional dissociation (HCD) fragmentation. Precursors were isolated using the quadrupole using a 1.6 m/z isolation width. An HCD collision energy of 25% was used, the AGC target was set to 2 × 10^5^ and an ion injection time of 50 ms was allowed. Dynamic exclusion of ions was implemented using a 45 s exclusion duration. An electrospray voltage of 1.8 kV and capillary temperature of 280°C, with no sheath and auxiliary gas flow, was used.

### Mass spectrometry data analysis

All spectra were analysed using MaxQuant 1.6.14.0 ^35^, and searched against the *Bacteroides thetaiotaomicron* Uniprot proteome database (UP000001414) downloaded on 22 September 2020. Peak list generation was performed within MaxQuant and searches were performed using default parameters and the built-in Andromeda search engine ^36^. The enzyme specificity was set to consider fully tryptic peptides, and two missed cleavages were allowed. Oxidation of methionine and N-terminal acetylation were set as variable modifications. Carbamidomethylation of cysteine was set as a fixed modification. A protein and peptide false discovery rate (FDR) of less than 1% was employed in MaxQuant. Proteins that contained similar peptides and that could not be differentiated on the basis of MS/MS analysis alone were grouped to satisfy the principles of parsimony. Reverse hits, contaminants, and proteins only identified by site modifications were removed before downstream analysis. Ranking of protein abundance was performed using iBAQ intensity values ^22^ obtained from MaxQuant. Proteomics data analysis was performed using the R package Limma ^37^. Paired Student’s t-tests were adjusted using a Benjamini-Hochberg and a FDR threshold of p-value <0.05 was applied.

### Construction of plasmids for protein expression in *E. coli*

The nucleotide sequences encoding Bt1760 (residues 2-503) and Bt1761 (residues 2-438) were amplified from B. theta genomic DNA, excluding the signal sequence and the lipid anchor cysteine. In all cases, the protein numbering starts with the first residue of the mature sequence, corresponding to C21 in Bt1760 and C24 in Bt1761 precursor amino acid sequences. The PCR product encoding Bt1760 was digested with NcoI and XhoI and ligated into pET28b, resulting in fusion of the coding sequence to a C-terminal LEHHHHHH tag. The PCR product encoding Bt1761 was digested with NdeI and XhoI and ligated into pET28a, resulting in fusion of the Bt1761 coding sequence to an N-terminal MGSSHHHHHHSSGLVPRGSHM tag. TOP10 cells were transformed with the ligation mixtures and plated on LB agar plates with kanamycin. After overnight incubation at 37°C, clones were screened for successful ligation by colony PCR. Bt1760 and Bt1761 variants were generated using the Q5 site directed mutagenesis kit (NEB). All constructs were verified by Sanger sequencing.

### Expression and purification of proteins from *E. coli*

Bt1760, Bt1761 and their variants were overexpressed in *E. coli*. Electrocompetent *E. coli* BL21(DE3) cells were transformed with the appropriate plasmid, plated on LB kanamycin plates and incubated at 37°C overnight. Starter LB kanamycin cultures were inoculated by scraping the transformants and incubated at 37°C, 180 rpm for 2 hours. 13 ml of the starter culture were used to inoculate each litre of LB kanamycin. The cultures were incubated at 37°C, 180 rpm until OD600 0.5-0.6 and induced with 0.2 mM IPTG. The temperature was then lowered to 20°C and the cultures were incubated for a further 19-21 h. The cells were harvested by centrifugation and the pellets were stored at −20°C.

The pellets were thawed, supplemented with DNase I and homogenised in TBS buffer. The cells were lysed with a single pass at 25 kpsi through a cell disruptor (Constant Systems). The lysates were supplemented with 1 mM PMSF. Unbroken cells were pelleted by centrifugation for 30 min at 30,000g, 4°C. The supernatants were loaded on a 5 ml Ni^2+^-charged chelating sepharose column by gravity flow at room temperature. The column was washed with 40 column volumes TBS buffer containing 30 mM imidazole, and the bound proteins were eluted with 5 column volumes TBS buffer containing 250 mM imidazole. The eluates were concentrated in an Amicon Ultra filtration device (30 kDa cut-off) by centrifugation. The samples were then loaded in batches on a HiLoad 16/600 Superdex 200 pg column (Cytiva) in 10 mM HEPES-NaOH pH 7.5, 100 mM NaCl. Elution fractions were collected and analysed by SDS-PAGE for purity. Fractions containing the proteins of interest were pooled, concentrated, flash-frozen in liquid nitrogen, and stored at −80°C.

### Fructooligosaccharide production

FOS used for cryo-EM, crystallography and ITC were generated by partial digestion of *Erwinia herbicola* levan (Sigma) by Bt1760 as described previously^12^.

### Isothermal titration calorimetry

Protein samples were thawed, centrifuged to remove any aggregates, and diluted to 25 or 50 μM in 10 mM HEPES-NaOH pH 7.5, 100 mM NaCl. Levan from *E. herbicola* or defined-length FOS were dissolved in the same buffer to 8 mg/ml and 1 mM, respectively. ITC was performed using a Microcal PEAQ-ITC instrument (Malvern Panalytical). Levan or FOS was injected into the sample cell containing protein or buffer. The titrations were performed at 25°C. The sample cell was stirred at 750 rpm. After an initial delay of 60 s, an injection of 0.4 μl was done (which was discarded from data analysis) followed by 18 injections of 2 μl. The spacing between injections was 150 s. Ligand to buffer control titrations were subtracted from all experiments. The experiments were repeated at least twice. Data were fitted to a single binding site model using the Microcal PEAQ-ITC Analysis software v1.40.

It was impossible to determine the molar concentration of the levan titrant due to its heterogeneity in chain length. Therefore, the molarity of available binding sites was estimated during data fitting. For Bt1760, the only secondary binding site substitution that had an effect on the affinity was W217A. We assumed that all the affinity observed for this variant could be attributed to binding to the active site alone. Therefore, the stoichiometry was fixed to 1 and the ligand concentration was floated during data fitting. The estimated molar concentration of 0.8% levan was 829 μM. N.B. the “molarity” in this case corresponds to the number of accessible binding sites for the enzyme on the polymeric levan ligand, rather than number of molecules, per volume. This titrant concentration was fixed for all other data fits for Bt1760. Similarly, by fixing n to 1, the levan titrant concentration for Bt1761 was determined to be 1.48 mM, suggesting ∼2x the number of binding sites on levan available to the SGBP as to the enzymes active site. Notably, the affinity of Bt1761 for levan determined this way was similar to that determined using defined-length FOS with known molarity (**ED figure 10**).

### Protein crystallisation

Bt1760_SeMet was crystalized in the presence of 200 mM potassium/sodium tartrate, 100 mM sodium citrate pH 5.6, 1.4 M ammonium sulphate and 500 mM fructose. Bt1760_D42N crystal forms were crystalized in 1.5 M lithium sulphate and 200 mM ammonium sulphate, 100 mM MES pH 6.5 and 30% PEG 5000 MME respectively. Bt1761 (both native and SeMet protein) was crystallised using 1.8-2.2 M (NH_4_)_2_SO_4_, 0.1 M MES pH 6.5. The protein concentrations were in the range of 10 mg/ml. The drops, composed of 0.1 µl or 0.2 µl of protein solution plus 0.1 µl of reservoir solution, were set up using a Mosquito crystallization robot (SPT Labtech). The vapor diffusion sitting drop method was used and the plates were incubated at 20 °C. If required, crystal hits were optimised via hanging drop vapour diffusion with larger volume drops (typically 1-1.5 µl). Bt1760_SeMet samples did not require additional cryoprotection. Bt1760_D42N samples were cryoprotected with paratone-N and with addition of 20% PEG 400 to the reservoir respectively. Bt1761 samples were cryoprotected by adding 4-fold excess of 3.5 M (NH_4_)_2_SO_4_ to the crystal drops.

### Data collection, structure solution, model building, refinement and validation

Diffraction data were collected at the synchrotron beamlines I02, I03 and I04 of Diamond Light Source (Didcot, UK) at a temperature of 100 K. The data set for Bt1760 SeMet was integrated with DIALS^38^ via XIA2^39^ and scaled with Aimless^40^. The space group was confirmed with Pointless^41^. The phase problem was solved by experimental phasing with Crank2^42^. Mutant BT1762_D42N data sets were integrated by XDS^43^ and processed as above. After phase transfer from experimental phasing the automated model building program task CCP4build on CCP4cloud^44^ delivered models with Rfactors below 30 %. The models were refined with Refmac^45^ and manual model building with COOT^46^. The final models were validated with COOT and MolProbity. Data collection and refinement statistics are presented in Supplementary Table 1. Other software used were from CCP4 suite^47^. Data collected for SeMet Bt1761 allowed solving the phase problem and partial model building via single anomalous dispersion (Se-SAD) using Phenix AUTOSOL^48^. Iterative rounds of manual building within COOT and model refinement in Phenix resulted in a partial model with R_free_ ∼35%, which was used as the input for model completion in the cryo-EM maps of the levan utilisome with long FOS. The segments missing from the Bt1761 X-ray model could not be modelled using the complete cryo-EM structure due to the poor quality of the X-ray electron density maps in the missing regions.

### Levan utilisome cryo-EM sample preparation, data collection and image processing

A sample of the purified apo levan utilisome complex (Bt1760-Bt1763) solubilised in DDM-containing buffer (10 mM HEPES, pH 7.5, 100 mM NaCl, 0.03% DDM) was prepared at 3 mg/ml. Lacy carbon 300-mesh copper grids (Agar Scientific) were glow-discharged in air (10 mA, 30s, Cressington 208). A sample volume of 3.5 mL was applied to the grid. Blotting and plunge freezing into liquid nitrogen-cooled liquid ethane were carried out using an FEI Vitrobot Mark IV (Thermo Fisher Scientific) with chamber conditions set at a temperature of 4 °C and 100% relative humidity. The grid was blotted for 6 s with a blot force of 6. Micrograph movies were collected on a Titan Krios Microscope (Thermo Fisher Scientific) operating at 300 kV with a Falcon III direct electron detector operating in counting mode. Data acquisition parameters can be found in Supplementary Table 2.

An initial dataset comprising 2057 micrograph movies was collected and image processing was carried out in Relion3.1^49^. Drift correction and dose-weighting was carried out using MOTIONCOR2^50^. CTF estimation of motion corrected micrographs was performed using Gctf^51^. Template-based particle picking within Relion was hindered by the large amount of carbon present in many micrographs. The micrograph stack was therefore manually culled to remove micrographs containing >50% carbon, leaving 1093 micrographs for further processing. Final particle picking was performed using the crYOLO general model^52^, yielding 96,639 particles. This particle stack was subjected to several rounds of 2D classification, after which 89,305 particles remained. A 3D starting model was generated *de novo* from the data using the stochastic gradient descent algorithm within RELION. 3D classification was used to isolate particles corresponding to the complete octameric utilisome complex (45,594). These particles were subjected to further rounds of classification in 3D to assess the conformational heterogeneity of the complex. Classification revealed considerable heterogeneity in the position of the SusD lids, with positions described as ‘wide open’ (W) and ‘narrow open’ (N) identified in all possible combinations. WW, WN and NN states contained 16,155, 22,452 and 6,987 particles respectively. To increase particle numbers and improve the results of downstream processing a second dataset of 3142 movies was collected. These were processed in the same way as described for the initial dataset, with particles picked using crYOLO. Classification in 3D yielded 146,512 particles that corresponded to the complete octameric utilisome complex. To improve the resolution for the more static, C2 symmetric portions of the utilisome (SusC, levanase and N-terminal region of the levan-binding protein subunits) particles stacks corresponding to the complete octameric complex from both datasets (192,106 total) were combined and subject to focused refinement with C2 symmetry. The mask applied in focused refinement excluded the SusD subunits. Particles were subjected to iterative rounds of CTF-refinement and Bayesian polishing (run separately for each dataset) until no further improvement in resolution was seen. Post-processing was performed using a soft, extended mask and yielded a global sharpened reconstruction at 3.5 Å, as estimated by the gold standard Fourier shell correlation using the 0.143 criterion.

### Active levan utilisome in the presence of FOS (DP8-12) cryo-EM sample preparation, data collection and image processing

A sample of the levan utilisome containing an active levanase solubilised in a DDM-containing buffer (10 mM HEPES, pH 7.5, 100 mM NaCl, 0.03% DDM) was prepared at 3 mg/ml and incubated with 0.5 mM levan FOS with a degree of polymerisation of ∼8-12 for at least one hour before grid preparation. Quantifoil carbon grids (R1.2/1.3, 300 mesh) were glow discharged (30 mA, 60 s, Quorum GloQube) in the presence of amylamine vapour. A sample volume of 3.5 μL was applied to the grid. Blotting and plunge-freezing into liquid nitrogen-cooled liquid ethane were carried out using an FEI Vitrobot Mark IV (Thermo Fisher Scientific) with chamber conditions set at a temperature of 4 °C and 100% relative humidity. The grid was blotted for 6 s with a blot force of 6. Micrograph movies were collected on a Titan Krios Microscope (Thermo Fisher Scientific) operating at 300 kV with a Falcon III direct electron detector operating in counting mode. Data acquisition parameters can be found in Supplementary Table 3.

A dataset comprising 974 micrograph movies was collected and image processing was carried out in RELION 3.1 ^49^. Drift correction and dose-weighting was done using MOTIONCOR2 ^50^. CTF estimation of motion corrected micrographs was performed using Gctf ^51^. Particle picking was performed using the general model of crYOLO and yielded 72,373 particles ^52^. Unwanted particles/contamination were removed from the particle stack through two rounds of 2D classification, after which 63,789 particles remained. Classification in 3D was used to address compositional heterogeneity. Good classes containing unambiguous SusC_2_D_2_ density represented the complete octameric utilisome, a hexameric assembly which lacked one levanase and one LBP subunit and a naked SusC_2_D_2_ complex. Contributing particles numbers were 17,045, 31,789 and 7390, respectively. SusD lids invariantly occupied a closed position and conformational heterogeneity was limited to the position of the levan binding protein.

Complete utilisome particles were subjected to iterative rounds of CTF-refinement and Bayesian polishing until no further improvement in resolution was seen. Post-processing was performed using a soft, extended mask and yielded a global sharpened reconstruction at 3.2 Å, as estimated by the gold standard Fourier shell correlation using the 0.143 criterion.

To extract higher resolution information for the SusC_2_D_2_ core complex, particle subtraction was performed to remove signal for additional lipoprotein components from all experimental images contributing to good classes (as defined above). A soft, expanded mask encompassing only the SusC_2_D_2_ core was generated using the volume eraser tool within Chimera ^53^ before using the resulting carved volume as in input for mask creation in RELION. Subtracted particles were used in a focused refinement with the same mask applied while enforcing C2 symmetry. Iterative rounds of CTF-refinement and Bayesian polishing were employed until no further improvement in resolution was observed. Post-processing resulted in a final sharpened reconstruction at 2.9 Å.

### Inactive levan utilisome in the presence of FOS (DP15-25) cryo-EM sample preparation, data collection and image processing

A sample of the levan utilisome containing inactivated levanase (D42A), solubilised in a DDM-containing buffer (10 mM HEPES, pH 7.5, 100 mM NaCl, 0.03% DDM), was prepared at 3 mg/ml and incubated with ∼0.5 mM levan FOS with a degree of polymerisation 15-25 for at least one hour before grid preparation. Grid type, preparation, microscope and detector were the same as for the active levan utilisome described above. Data acquisition parameters can be found in Supplementary Table 4.

A dataset comprising 1388 micrographs movies was collected and image processing was carried out in RELION 3.1 ^49^. Drift correction and dose-weighting were carried out using MOTIONCOR2 ^50^. Particle picking was performed using the general model of crYOLO and yielded 157,953 particles^52^. Unwanted particles/contamination were removed from the particle stack through two rounds of 2D classification, after which 146,056 particles remained. Classification in 3D was used to address compositional heterogeneity. Good classes representing the complete octameric utilisome and the hexameric assembly lacking one levan binding protein and one levanase subunit were observed and contained 78,469 and 42,488 particles, respectively. Conformational heterogeneity in the position of the levan binding protein was observed with some classes possessing a conformation where this subunit was held close to the levanase. Particles contributing to all classes with evidence of this docked conformation of the levan binding protein were pooled (98,755) and a consensus refinement was carried out. CTF refinement and Bayesian polishing were performed iteratively until no further improvement in resolution was observed. The resulting reconstruction possessed weaker density for the C-terminal domain of the levan binding protein than for the N-terminal portions. To improve this, a focused classification approach without alignment was used.

A mask encompassing only the docked position of the levan binding protein with some surrounding density was created using the volume eraser tool in Chimera followed by mask creation in RELION. A focused 3D classification job without alignment was run using the aforementioned mask and the output from the aforementioned refinement as a reference model. The reference model was low-pass filtered to 3.5 Å, just above the resolution of 3.3 Å reported for the consensus refinement, thus allowing classification on high resolution features. Several T values were empirically tested and a T value of 40 was found to give the best results. A single class, containing 27,310 particles, was identified that possessed well resolved density for the C-terminal domain of the levan binding protein. A particle star file containing information for particles contributing to this class was created manually via command line arguments. From this, two new star files were generated that contained random half sets of the selected data. Using relion_reconstruct, these star files were used to generate two independent half maps that corresponded to the unmasked structure. Post-processing using these generated half maps yielded a sharpened reconstruction of 3.0 Å, as estimated by gold standard Fourier Shell correlations using the 0.143 criterion. The density for the C-terminal domain of the levan binding protein was improved, and density corresponding to levan chain that links the levan binding protein to the levanase was also visible.

To obtain the highest quality density for FOS molecules occupying the SusCD binding cavity, all SusC_2_D_2_-containing particles were considered regardless of lipoprotein complement or conformation. A particle subtraction and focused refinement strategy targeting the SusC_2_D_2_ core of the complex was used as described for the active levan utilisome. Post-processing of the model arising from this final C2 symmetrised refinement resulted in a sharpened reconstruction at 2.7 Å.

### Dextran utilisome cryo-EM sample preparation, data collection and image processing

A sample of the dextran utilisome complex (Bt3087-Bt3090) solubilised in a DDM-containing buffer (10 mM HEPES, 100 mM NaCl, pH 7.5, 0.03 % DDM) was prepared at 0.05 mg/ml. Lacy carbon 300-mesh copper grids coated with a <3 nm continuous carbon film (Agar Scientific) were glow-discharged in air (10 mA, 30 s). A sample volume of 3.5 μL was applied to the grid. Blotting and plunge-freezing were performed 10 seconds after loading the sample onto the grid using an FEI Vitrobot Mark IV (Thermo Fisher Scientific) with chamber conditions set at a temperature of 4 °C and 100% relative humidity. The grid was blotted for 6 s with a blot force of 0. Micrograph movies were collected on a Titan Krios Microscope (Thermo Fisher Scientific) operating at 300 kV with a Falcon IV direct electron detector operating in counting mode. Data acquisition parameters can be found in Supplementary Table 5.

A dataset comprising 6,331 micrographs was collected and image processing was carried out in RELION 3.1 ^49^. Drift correction and dose-weighting were performed using RELION’s own implementation of MOTIONCOR2. CTF estimation of motion corrected micrographs was done using CTFFIND4 ^54^. Particle picking was done using the crYOLO general model which identified 820,184 particles in the micrographs. Junk particles and contaminants were removed through several rounds of 2D classification, after which 477,707 particles remained. An initial model was generated *de novo* from the data. Extensive 3D classification was used to address the considerable compositional heterogeneity that was present in the data (see **ED Figure 5**). Each unique composition was refined and sharpened independently. A consensus refinement was carried out, with iterative rounds of CTF-refinement and Bayesian polishing until no further improvement in resolution was seen. A final, sharpened consensus reconstruction was obtained at 3.1 Å.

### Model building into electron microscopy maps

Buccaneer (part of CCP-EM v1.5.0)^55, 56^ was used to build the initial protein models into the post-processed consensus inactive levanase map, resulting in almost complete protein models. An AlphaFold 2^23^ prediction of the dextran SusC was generated and used as an initial model. Manual modelling and real space refinement of protein and FOS chains were performed iteratively using COOT^46^ and Phenix^48^, respectively. The completed protein and FOS models were placed into other maps and real space refined. The acyl-cysteine was designated as the first residue of each lipoprotein. Model refinement statistics are presented in Supplementary Tables 2-5.

### Density analysis and figure making

Investigation and comparison of EM density maps was performed using Chimera^53^ and COOT^46^. Figures of maps and models were generated using Chimera and ChimeraX^57^. To aid interpretability of EM density in generated figures, some maps were filtered using LAFTER^27^. Maps processed in this way are clearly indicated in the corresponding figure legend. The masks supplied in filtering were the same masks used for post-processing within RELION.

## Extended Data

**Extended Data Figure 1.**
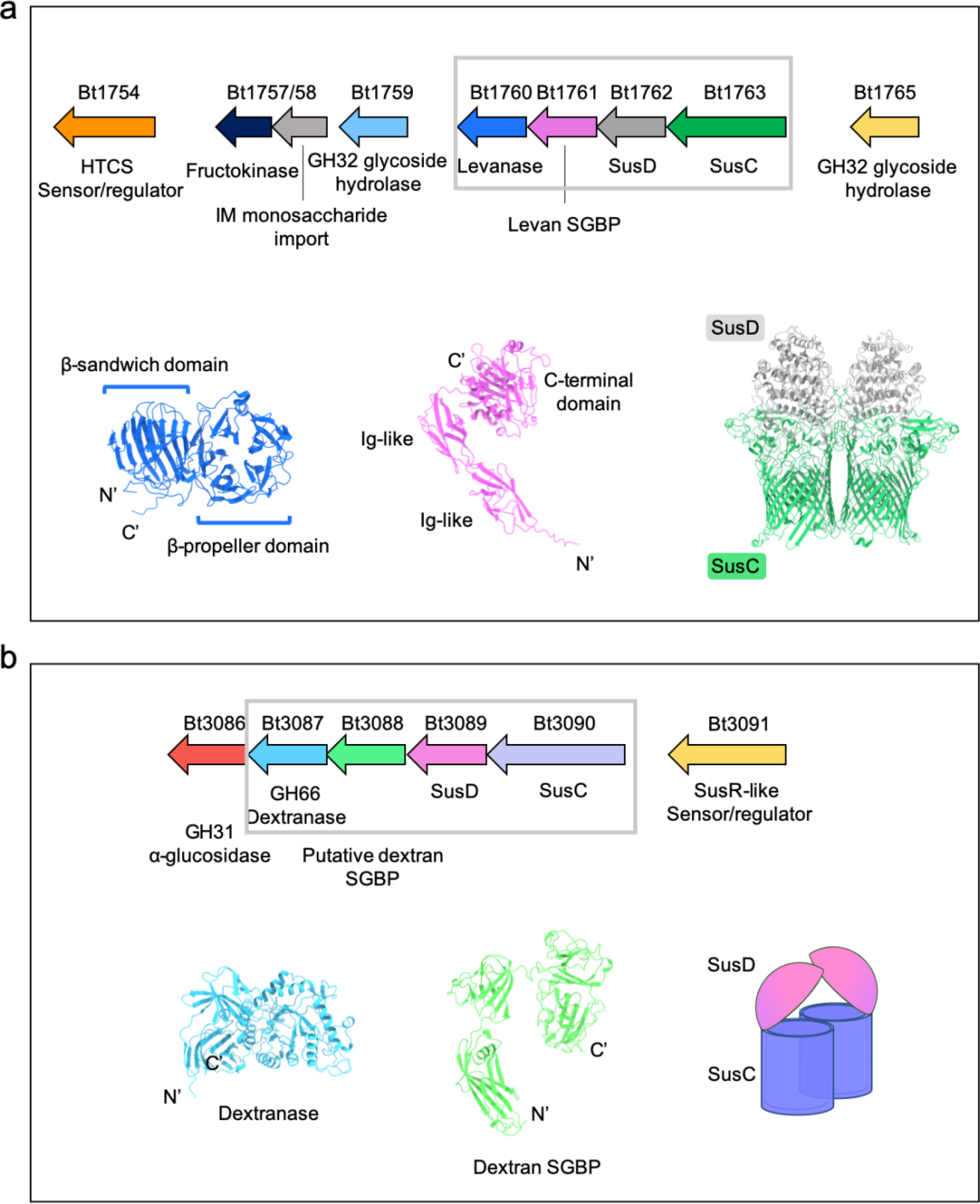
Arrangement of levan and dextran PULs with known and predicted structural information for OM-associated components. **a,** Organisation of the levan PUL showing gene positions within the locus with functions labelled. OM-associated PUL components are within the grey box. An X-ray crystal structure of the levanase Bt1760 is shown (blue) with its β-sandwich and catalytic β-propeller domains clearly identifiable (PDB ID 6R3R). The AlphaFold2-predicted model for Bt1761 is shown (pink) oriented such that the N-terminus is at the bottom and the globular C-terminal domain hypothesised to bind levan is at the top. Note that the N-termini of the additional lipoproteins will be lipidated and associated with the OM. The cryo-EM structure of the dimeric SusCD complex in the open-open state is shown with SusC components in green and SusD components in grey. **b**, Organisation of the dextran PUL showing gene positions within the locus with functions labelled. OM-associated PUL components are boxed in grey. AlphaFold2-predicted models for the dextranase (Bt3087) and putative dextran SGBP (Bt3088) are shown in blue and green, respectively.

**Extended Data Figure 2.**
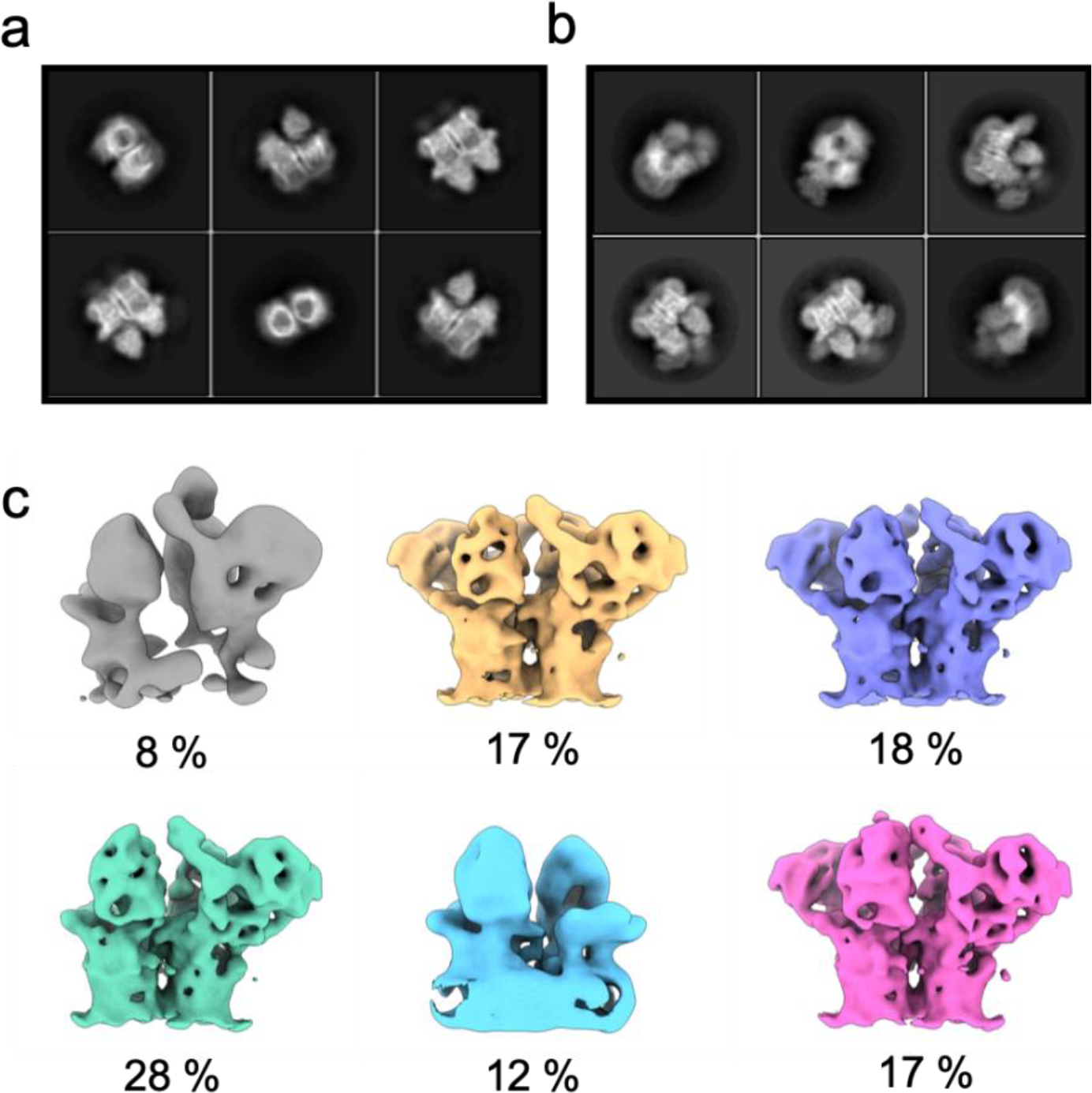
Early cryo-EM processing outputs of the apo levan SusC_2_D_2_ core complex and apo levan utilisome. **a,** Example 2D class averages of the LDAO-solubilised levan core complex from our previous study. **b**, Example 2D class averages of the levan utilisome solubilised in DDM, and with the HIS-tag moved from SusD to the levan SGBP. Density in addition to that of the core complex (**a**) can be seen in all views. **c**, Output of the first round of 3D classification. Yellow, purple and pink classes represent the octameric complex i.e. the complete utilisome. The green class shows the additional lipoproteins associated with just one SusC unit whilst the blue class shows that a small proportion of SusC_2_D_2_ complexes were present in isolation.

**Extended Data Figure 3.**
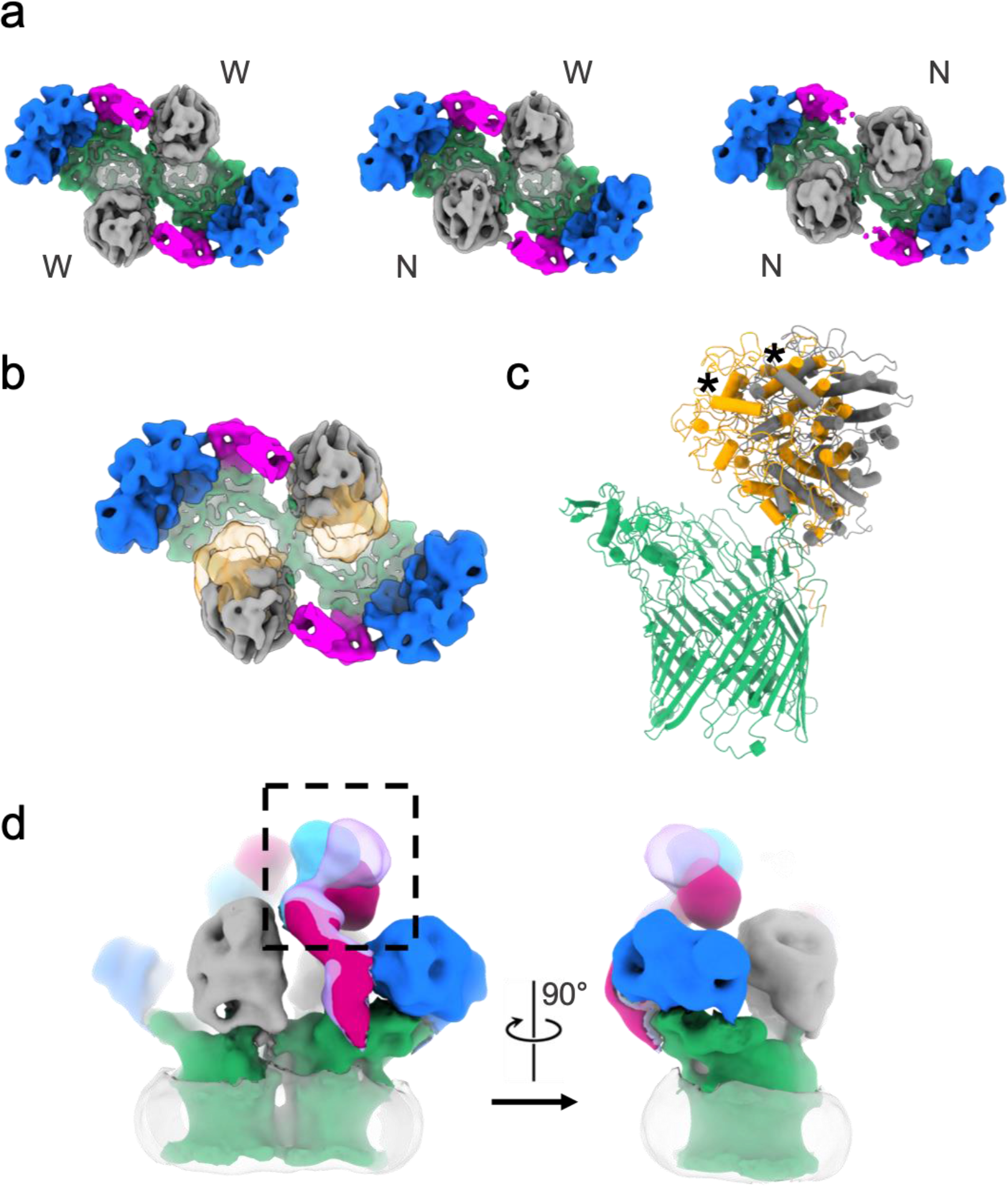
Conformational variability of the substrate-free levan utilisome. **a,** Refined outputs from 3D classification of the octameric utilisome viewed from the extracellular space. Bt1763 SusC (green), Bt1762 SusD (grey), Bt1761 levan SGBP (magenta), Bt1760 levanase (blue). Classes are separated on the basis of SusD lid position. Wide-wide, normal-wide, and normal-normal open states are presented from left to right with wide and normal positions labelled ‘W’ and ‘N’, respectively. **b**, Overlay of the wide-wide (grey) and normal-normal (orange) open states of the complex. **c**, Overlay of models for the normal open versus wide open state of the transporter generated by a rigid-body fit of SusD into the density of the respective maps. A monomer is shown for clarity and an asterisk marks the same SusD helix in both models. **d**, A view of the utilisome shown at high threshold levels in the plane of the membrane (left). Different conformations of the levan binding protein observed in 3D classification are overlaid to demonstrate the flexibility of this subunit (boxed region). The same view rotated 90° is shown (right). Disordered micelle density is shown as translucent grey.

**Extended Data Figure 4.**
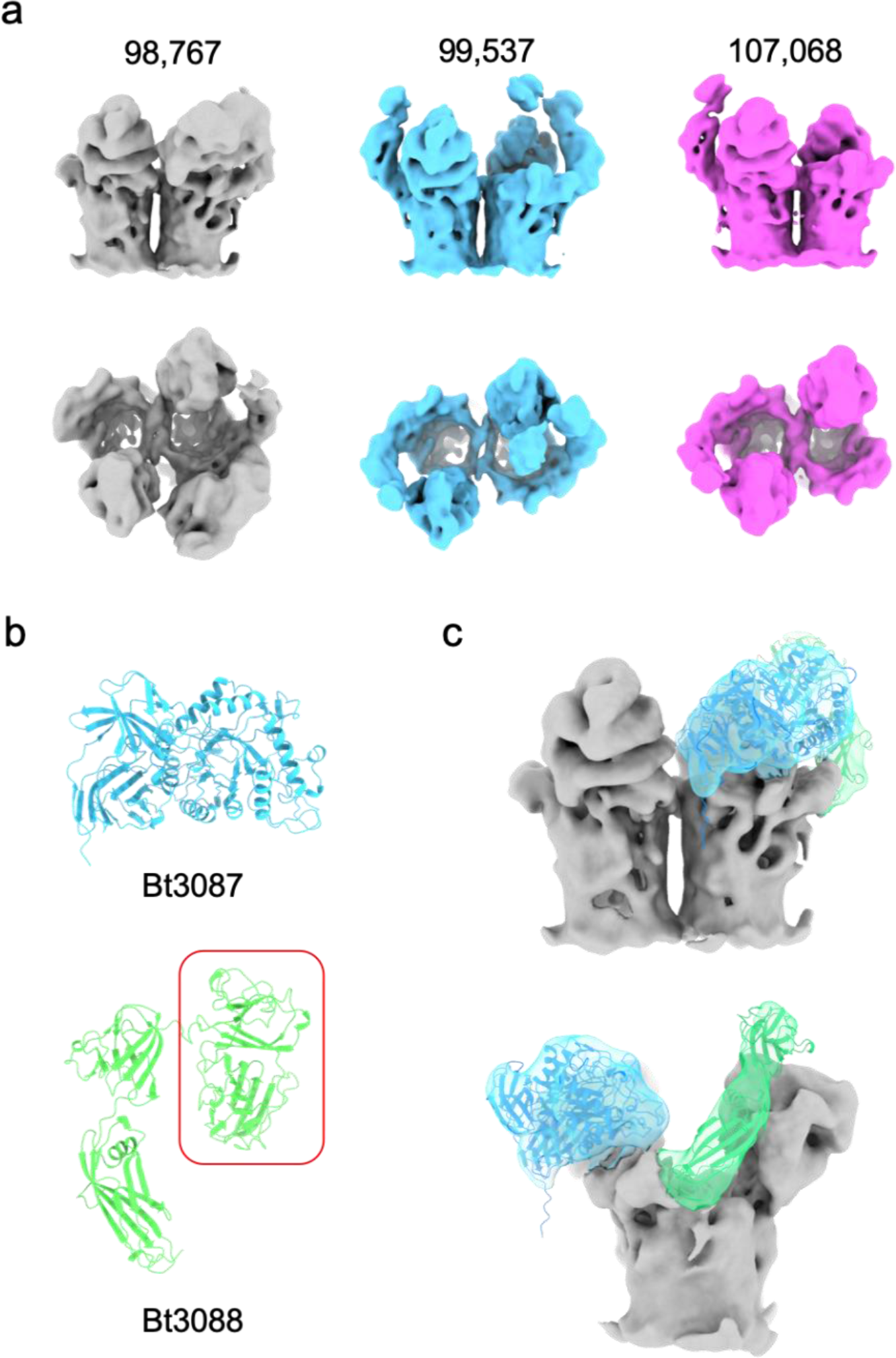
Initial outputs from 3D classification of the dextran utilisome and assignment of lipoprotein density. **a**, Good 3D classes with clear density for the core transporter viewed in the plane of the membrane (top) and from the extracellular space (bottom). Unassigned density was also present in all classes. Particle numbers contributing to each class are displayed. **b**, AlphaFold2 predictions of the dextranase (Bt3087; blue) and putative dextran SGBP (Bt3088; green) of the dextran utilisome. **c**, Grey class shown in (**a**) with additional densities assigned to the dextranase and putative SGBP on the basis of predicted structures. Note that the C-terminal domain of the SGBP (red box) is not visible in the EM map and the docked model for the SGBP has been truncated accordingly.

**Extended Data Figure 5.**
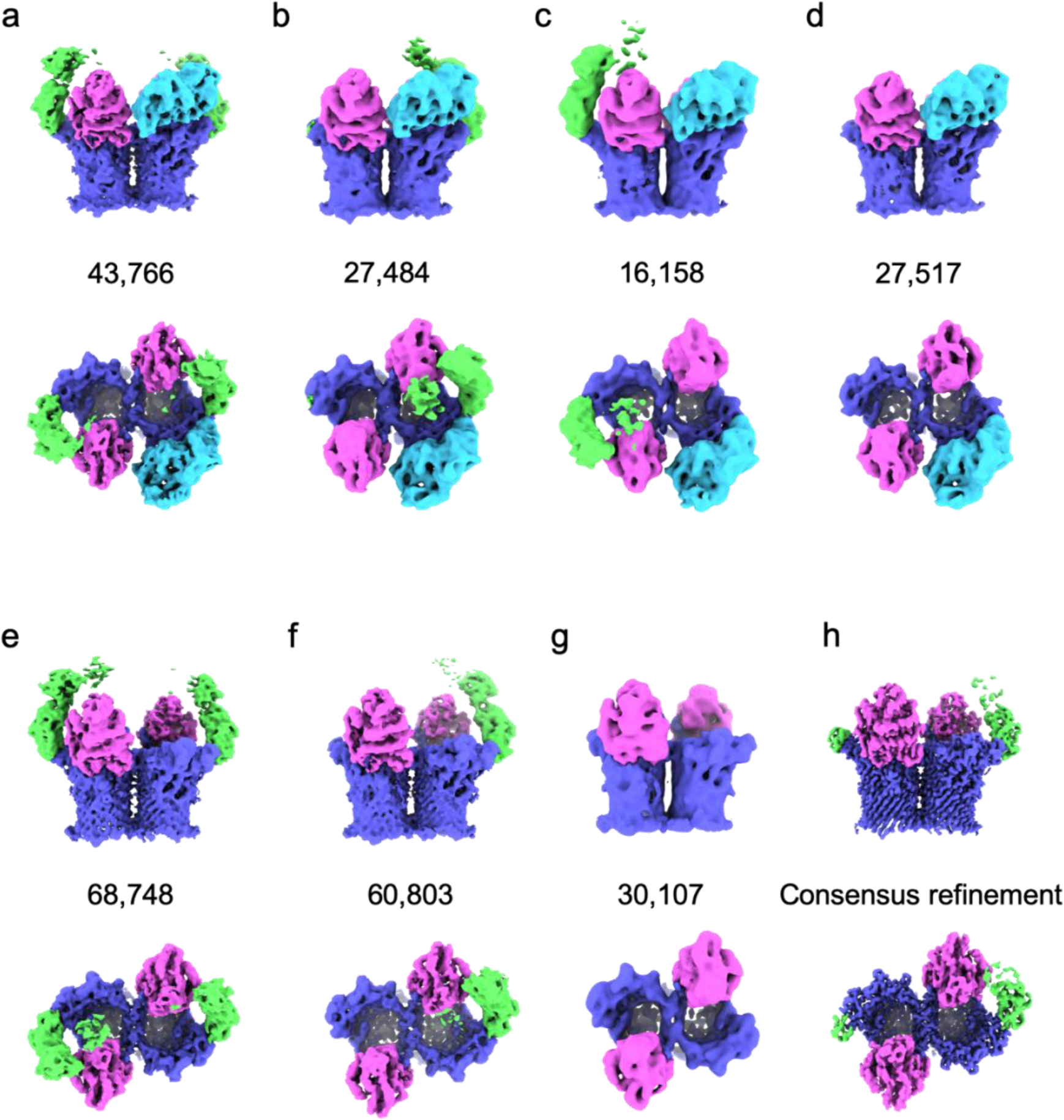
Compositional heterogeneity of the dextran utilisome observed by cryo-EM. **a-g**, Refined outputs of 3D classification viewed in the plain of the membrane (top panels) and from the extracellular space (bottom panels), where each map corresponds to a unique complement or arrangement of auxiliary components. SusC (Bt3090) is purple, SusD (Bt3089) is pink, the dextran SGBP (Bt3088) is green and the dextranase (Bt3087) is cyan. **h**, A consensus refinement at a global resolution of 3.1 Å.

**Extended Data Figure 6:**
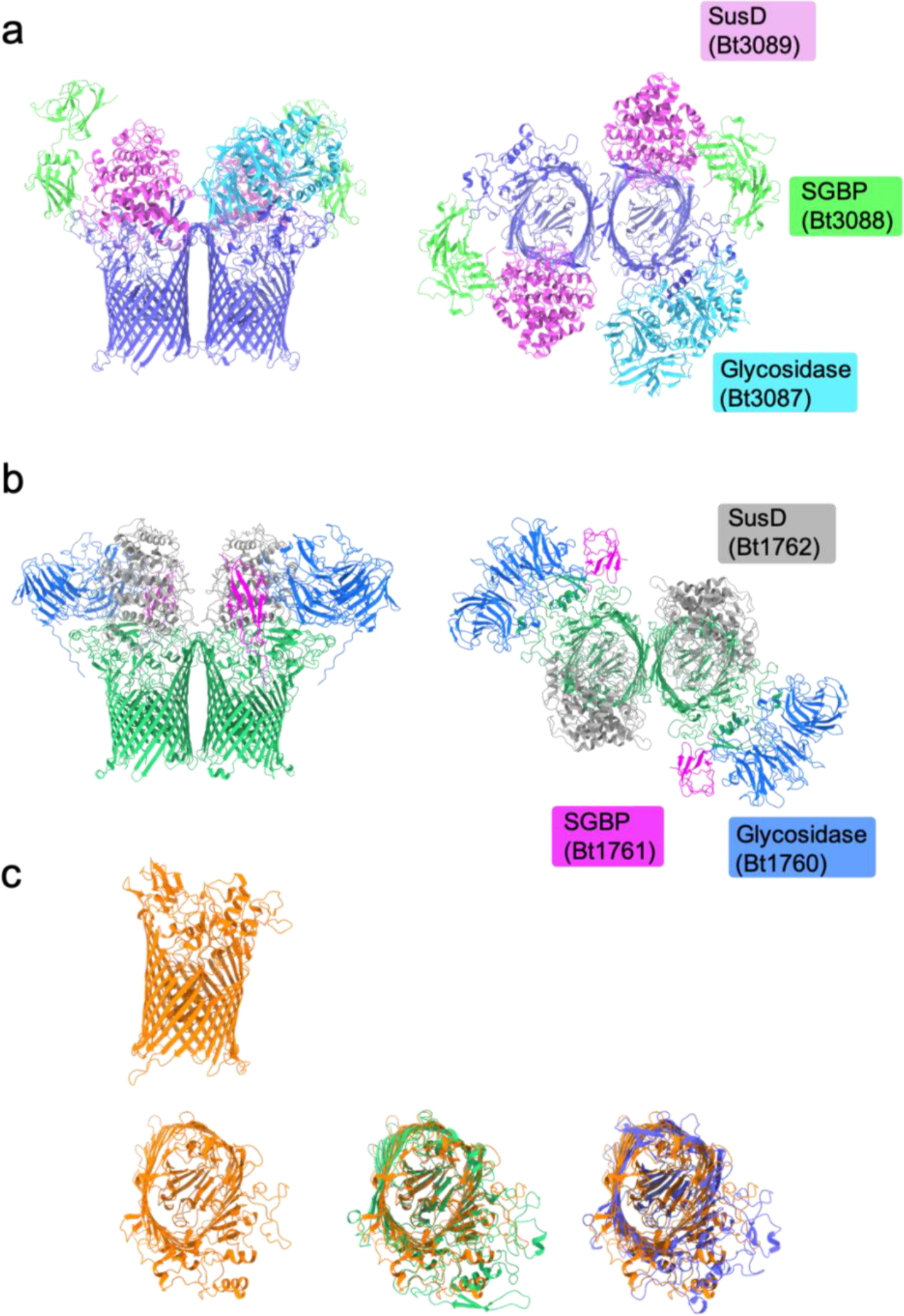
Arrangement of auxiliary proteins in apo glycan utilisomes. **a**, Composite model demonstrating the arrangement of the dextran utilisome. Cryo-EM data permitted refinement of the SusC components. AlphaFold2 structure predictions for SusD (Bt3089) and the dextranase (Bt3087) were obtained and docked into the cryo-EM map for the heptameric complex. An AlphaFold2 structure prediction for part of the SGBP (Bt3088) was also rigid fit to the cryo-EM density. Unambiguous density is visible only for the first two domains of the SGBP. The predicted model was therefore truncated prior to the C-terminal domain. **b**, Structure of the substrate-free levan utilisome. SusD components (Bt1762) are positioned based on a rigid body fit to a low-resolution map in which unambiguous density for the lids was observed. Note the different arrangement of the glycosidase and SGBP components relative to SusD in the levan and dextran systems. **c**, AlphaFold2 structure prediction of Bt3702 SusC from the starch utilisation system (orange) viewed in the plane of the membrane (top) and from the extracellular space (bottom). Alignments of the predicted structure with experimentally determined structures for SusC components of the levan (green) and dextran (violet) systems are shown.

**Extended Data Figure 7.**
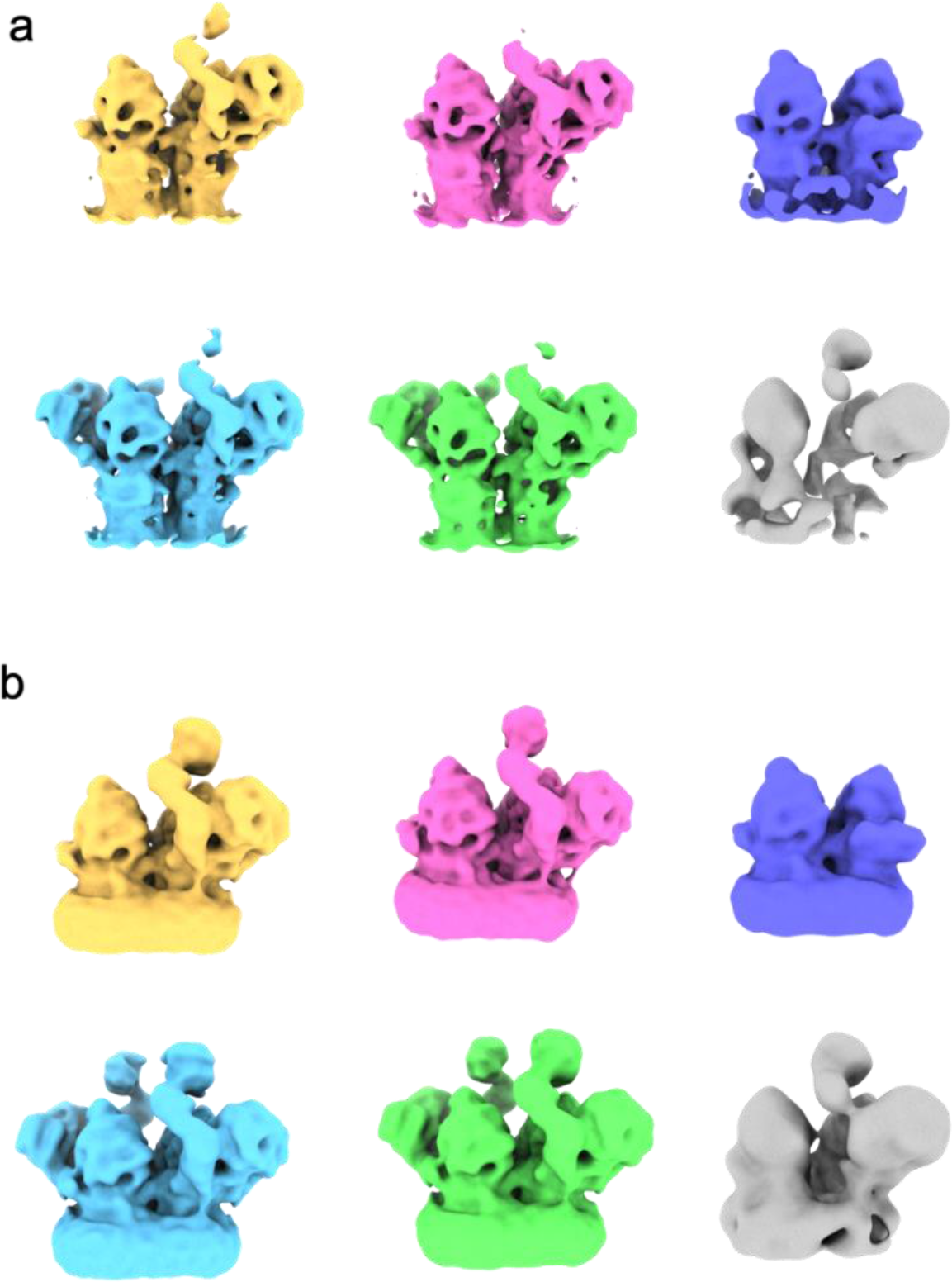
Output of 3D classification for the levan utilisome with an active levanase in the presence of FOS DP8-12. **a**, Classes viewed in the plane of the membrane. Classes containing particles of the complete octameric complex were observed (blue and green) as well as hexameric complexes containing just a single copy of the LBP and glycosidase (pink and yellow). A class containing SusC_2_D_2_ in isolation is also present (purple). **b**, The same classes as in (**a**) viewed at an increased contour level

**Extended Data Figure 8.**
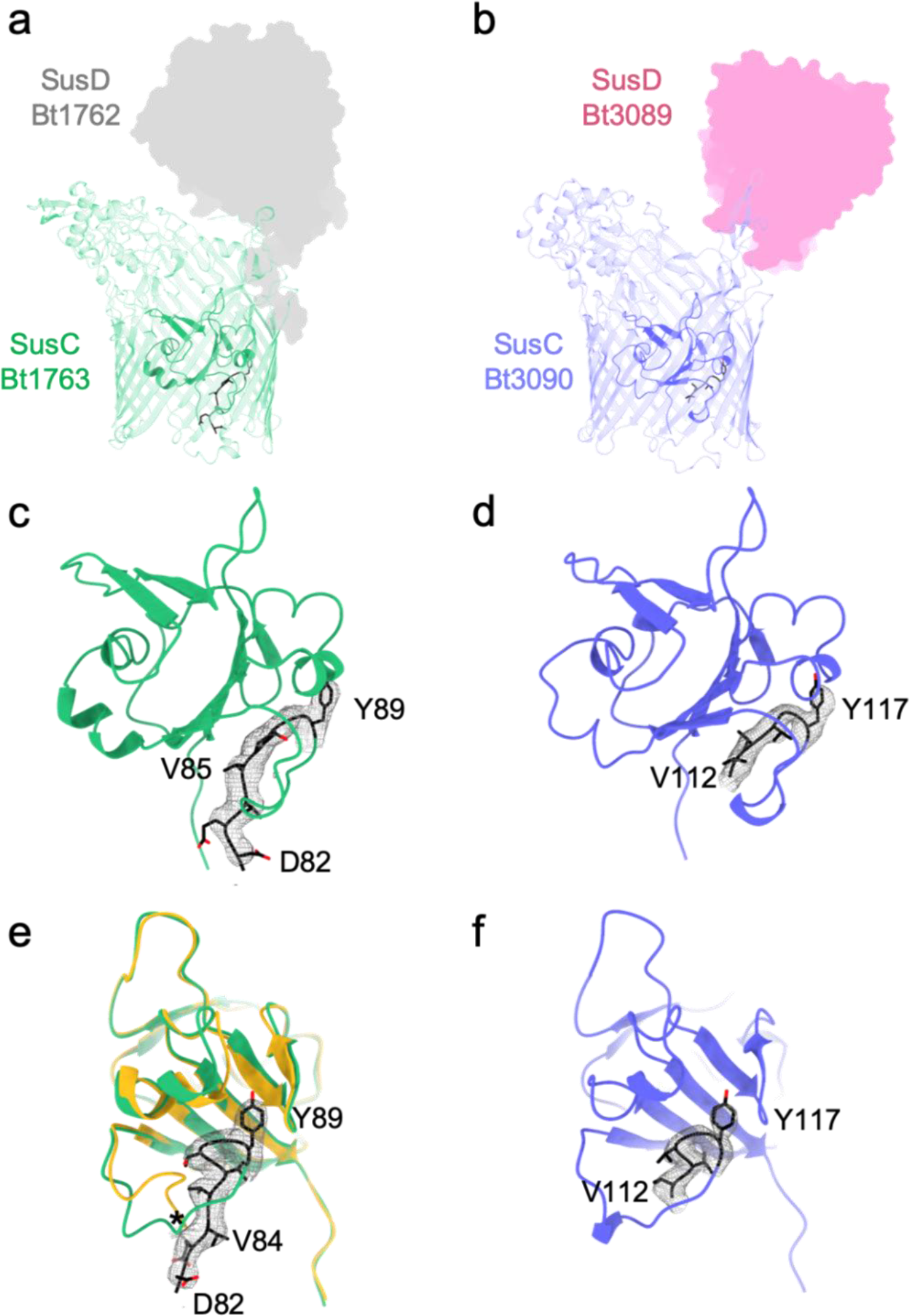
Analysis of the TonB box region in cryo-EM structures of the levan and dextran transporters. **a,** Structure of SusC of the levan transporter (Bt1763) in the absence of substrate. **b**, Structure of SusC of the dextran transporter (Bt3090) in the absence of substrate. Ton box regions are shown in black. The corresponding SusD components are shown as shaded zones; both occupy an open position. Isolated plug domains for each transporter can be seen in **c** (levan) and **d** (dextran). Density is displayed for the Ton box regions (residues in black stick form). **e**, Superposition of plug domains for the apo (green) and substrate-bound (yellow) structures of the Bt1763; rotated 90° relative to (**c**). The visible N-terminus is D82 for the substrate-free transporter compared with R93 for the substrate-bound transporter (marked with an asterisk). **f**, A view of the Bt3090 plug from the same angle as (**e**).

**Extended Data Figure 9.**
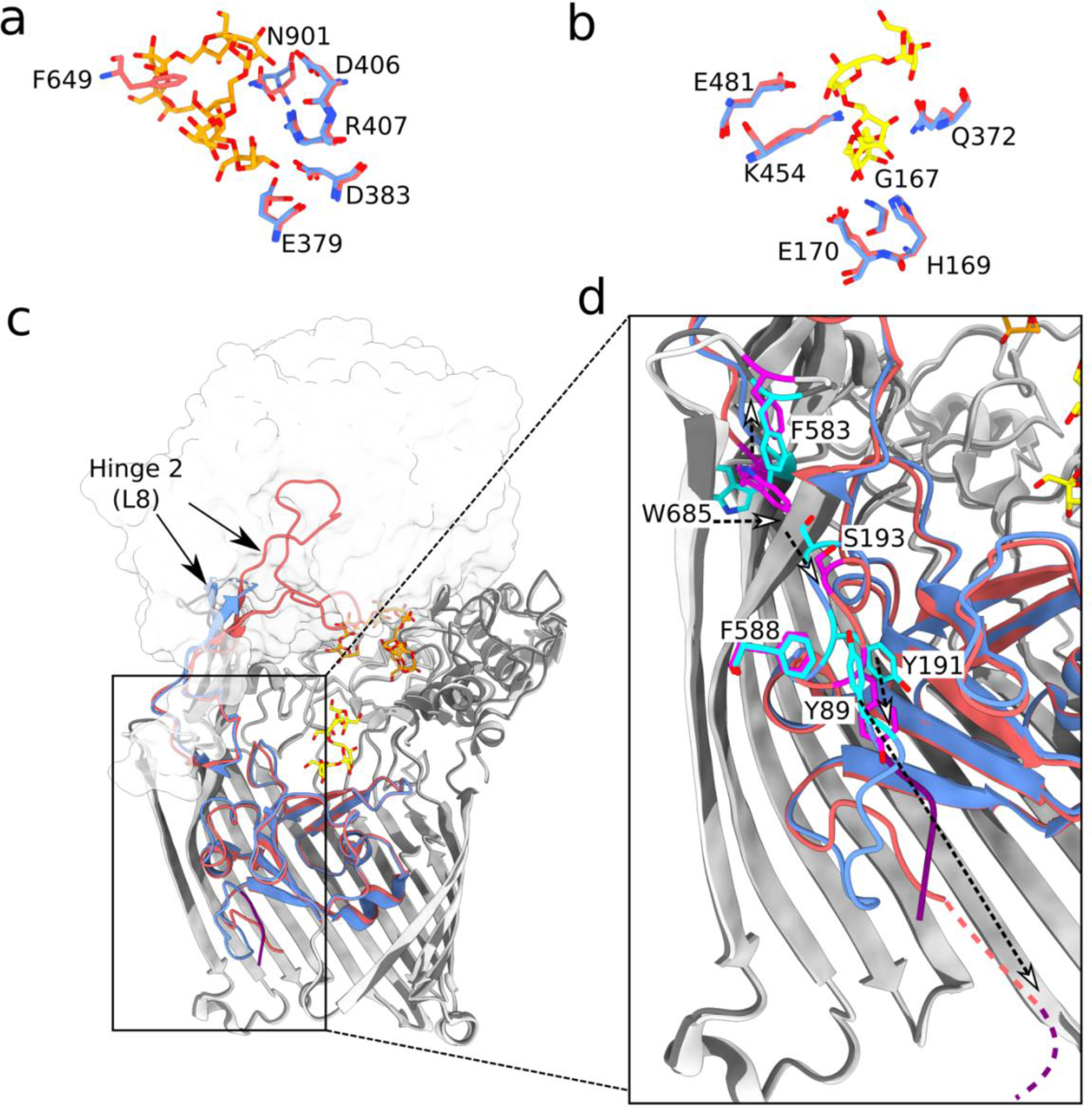
Subtle conformational changes upon substrate binding to Bt1763 SusC result in exposure of the Ton box. **a** and **b**, Overlay of apo (blue) and substrate-bound (salmon) Bt1763 FOS binding residues in the top (a) and bottom (b) binding sites. G167, H169 and E170 are part of the plug domain. **c**, Overlay of apo (dark grey) and FOS-bound (light grey) Bt1763 cryo-EM structures. Hinge 2 (L8^12^)and the plug domain in the apo and substrate-bound structures are coloured in blue and salmon, respectively. β-strands 13-22 of the SusC barrel are hidden. The closed SusD lid is represented as a transparent surface. **d**, a ∼90° rotated close-up view of the inset in **c**. Residues involved in transmitting substrate binding information from the extracellular side of the outer membrane to the periplasmic side are coloured in cyan and magenta in the apo and substrate-bound structures, respectively. The dashed white-headed arrows indicate residue movement from the apo to the substrate-bound state. The Ton box in the apo structure is shown in solid purple. The dashed line indicates the possible location of unresolved residues preceding R93 in the substrate-bound structure based on weak density observed in cryo-EM maps (Ton box in purple, residues 89-92 in salmon).

**Extended Data Figure 10.**
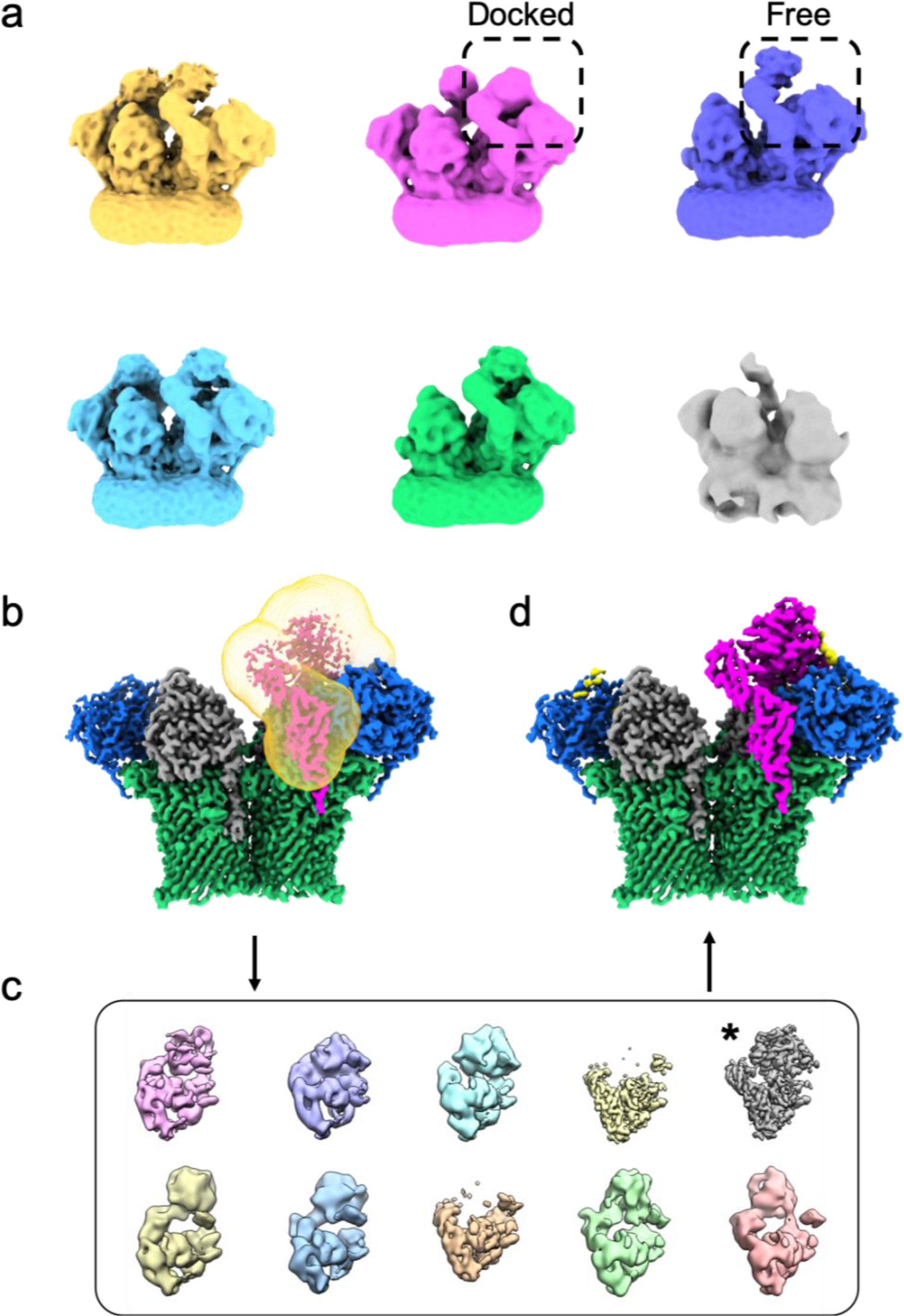
Addition of FOS DP ∼15-25 FOS to levan utilisomes containing an inactive levanase allows structure determination of the levan SGBP. **a**, Outputs of 3D classification showing that the levan SGBP can adopt a ‘docked’ conformation proximal to both the SusD and levanase. **b**, A consensus refinement of all classes containing at least one docked SGBP (yellow, pink, cyan and green). A mask was created around the region of interest (transparent yellow). **c**, Outputs of focused classification on the masked region without alignment. A class displaying high resolution for the region of interest is marked with an asterisk. Independent half maps were reconstructed using particles belonging to this class. **d**, Sharpened reconstruction generated with the aforementioned half maps showing improved density for the levan SGBP. Note that the handedness of maps displayed here is incorrect. This was addressed prior to model building.

**Extended Data Figure 11.**
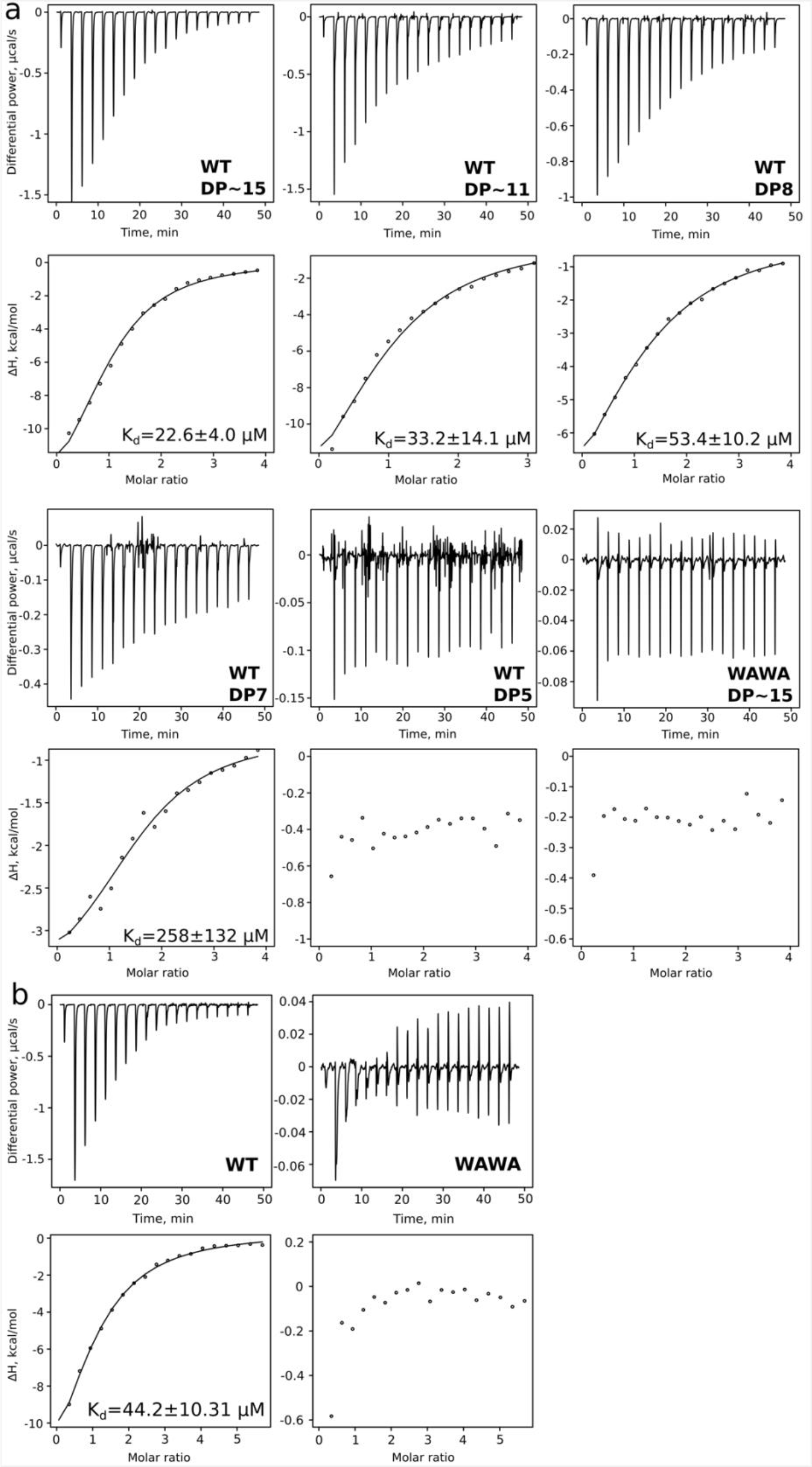
ITC with recombinant Bt1761 levan SGBP demonstrates importance of Trp297 and Trp359 for levan binding. **a**, Titration of 1 mM defined-length FOS into 50 μM wild type or W297A/W359A double-mutant (WAWA) Bt1761. The traces with FOS DP∼15 are the same ones shown in Figure 5e, and are repeated here for comparison with FOS of different sizes. The minimal FOS size for reasonably strong binding is DP 8. **b,** Titration of 8 mg/ml levan into 25 μM Bt1761. The affinity estimates, where possible, were obtained by fitting the data to a single binding site model (Methods).

**Extended Data Figure 12.**
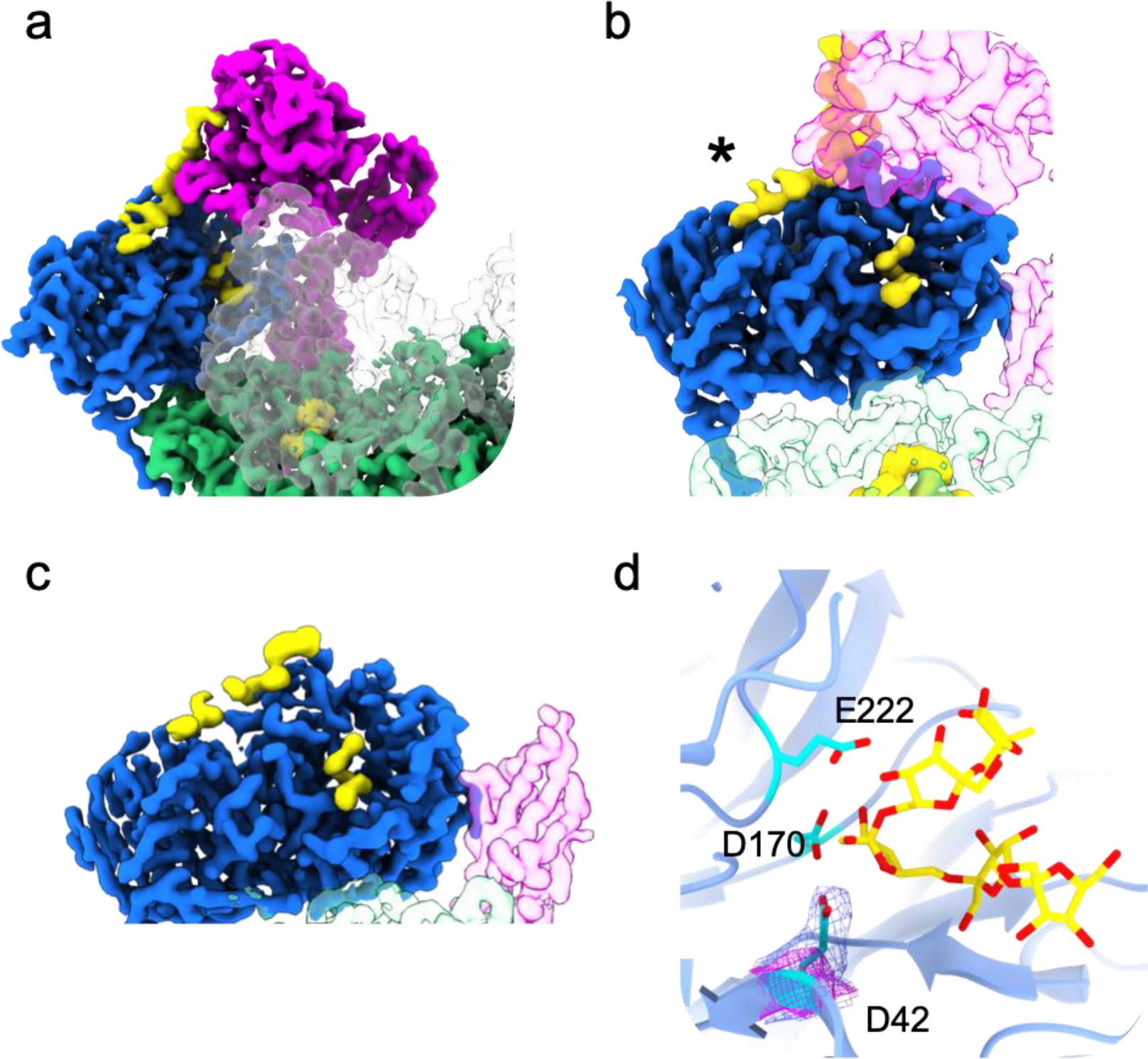
FOS binds to two distinct sites on the levanase. **a**, Zoomed view of one half of the levan utilisome highlighting the tethered position of the levan SGBP relative to the levanase, the SusD subunit (translucent density) and the top of the SusC barrel (green). **b**, Face on view of the levanase showing the distinct densities for levan (yellow) bound at the secondary site (marked with an asterisk) and at the active site. No density linking these two sites was observed. In both (**a**) and (**b**) FOS bound at the upper site in SusC is observed. The minimum distance between FOS at the active site of the levanase and in the binding cavity of SusC is ∼30 Å. **c**, View of the levanase associated with the alternate SusC unit where no tethering of the SGBP is observed but density for levan is still evident, bound across the top of the β-propeller and β-sandwich domains of the levanase. **d**, Zoomed view showing levan FOS modelled into the active site of the inactive levanase. Residues comprising the catalytic triad^28^ are shown as stick models. The inactive levanase possesses a D42A mutation which is clearly resolved when overlaying density from the active (blue) and inactivated (pink) levanase reconstructions.

**Extended Data Figure 13.**
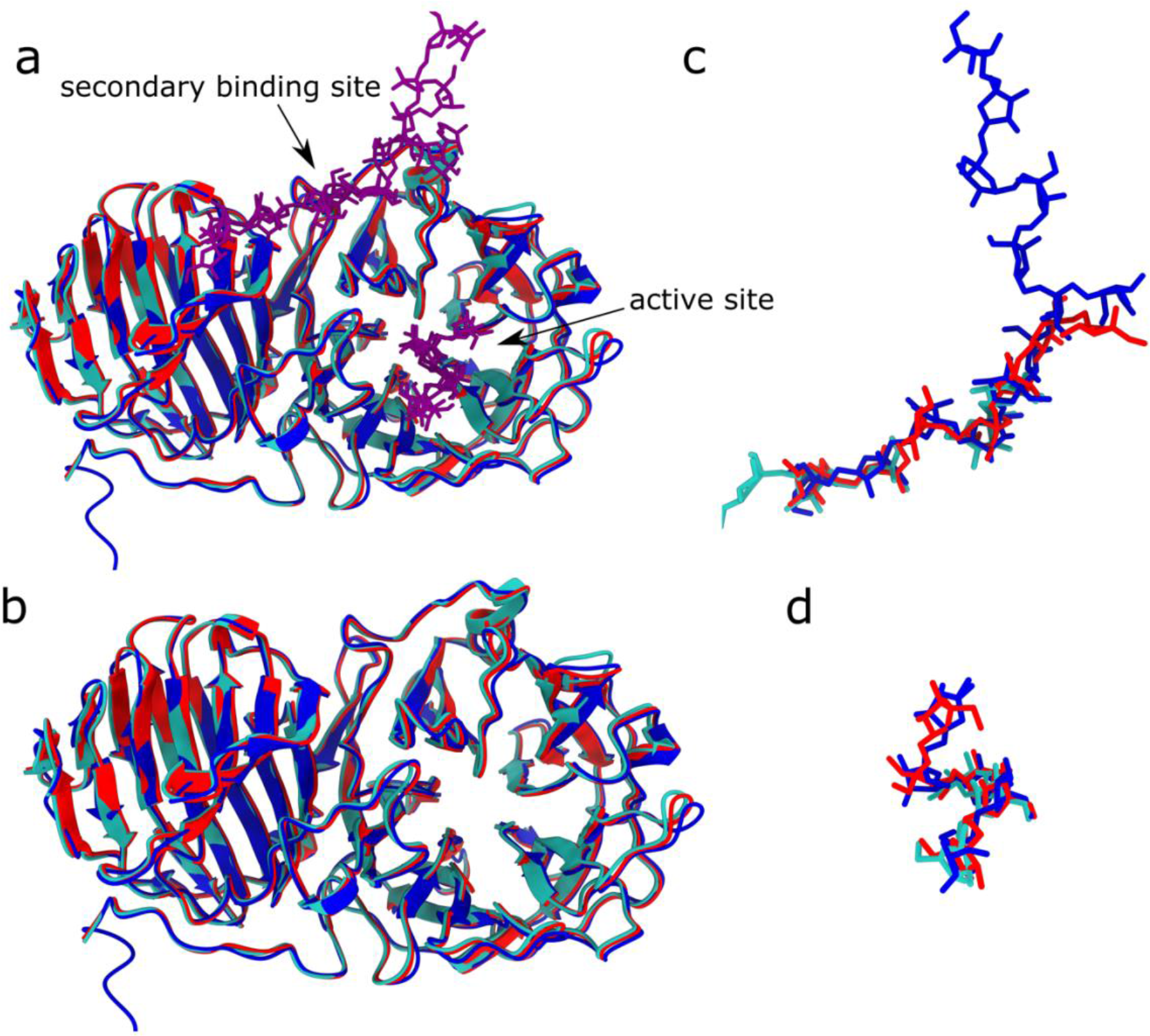
FOS-bound levanase structures determined by cryo-EM and X-ray crystallography are very similar. **a, b,** Cartoon representation of structures aligned using ChimeraX Matchmaker: the cryo-EM structure from the inactive levanase complex (blue), and two crystal structures of the levanase alone (red, cyan; 7ZNR and 7ZNS), with the FOS from all structures coloured in purple (not shown in **b**). **c** and **d**, Alignment of FOS in the secondary binding site and the active site, respectively. Colours are the same as the protein chains in **a** and **b.**

**Extended Data Figure 14.**
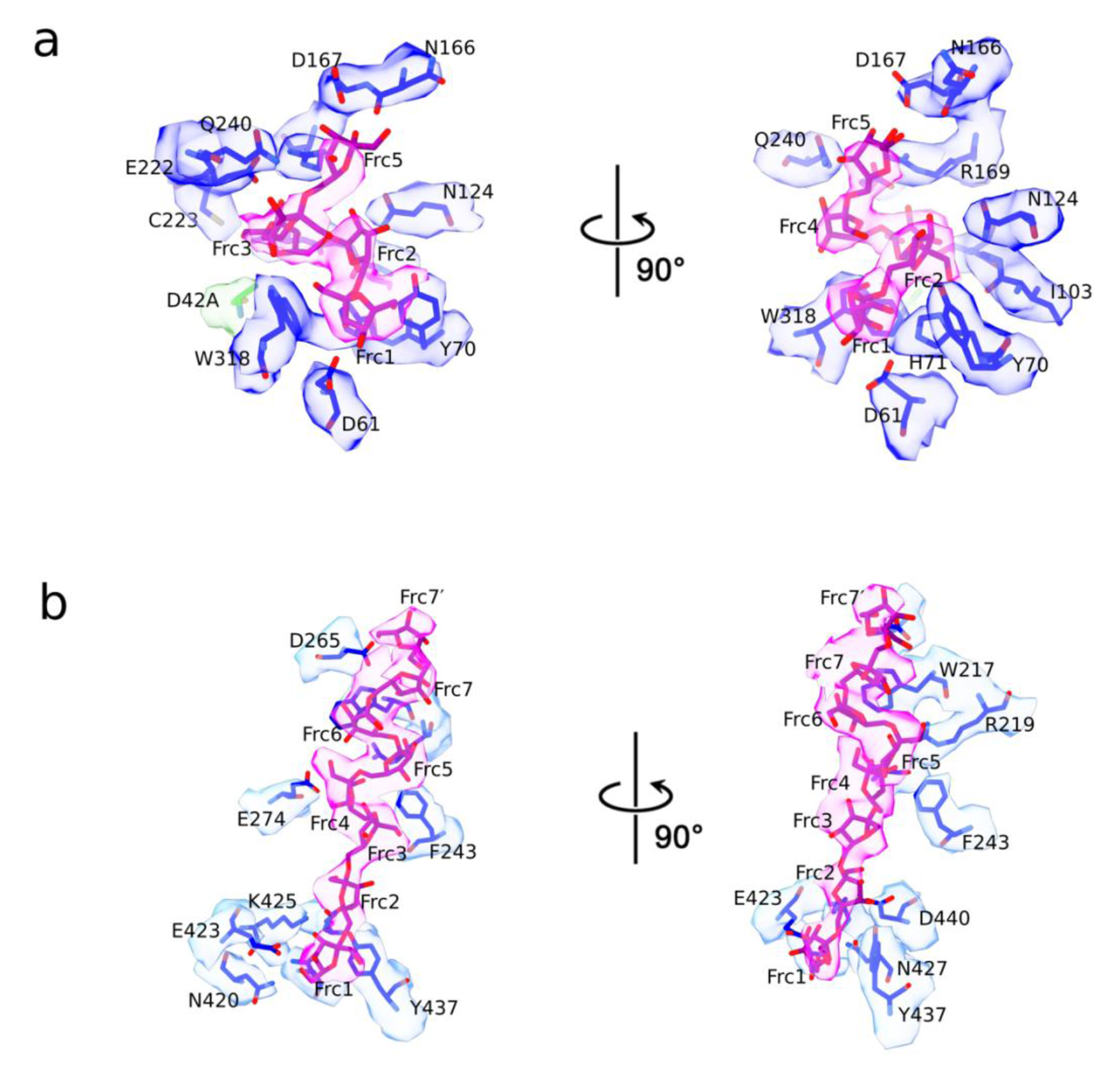
The cryo-EM structure shows that the interaction network between FOS and the levanase is extensive. Residues likely interacting with FOS in the active site of Bt1760 levanase, **a**, and in the secondary binding site, **b,** shown in blue. The FOS is displayed in magenta. The active site mutation, D42A, is shown in green. Frc7′ denotes the β2,1 decoration on Frc7. The EM density from the inactive levanase reconstruction is shown around each residue.

**Extended Data Figure 15.**
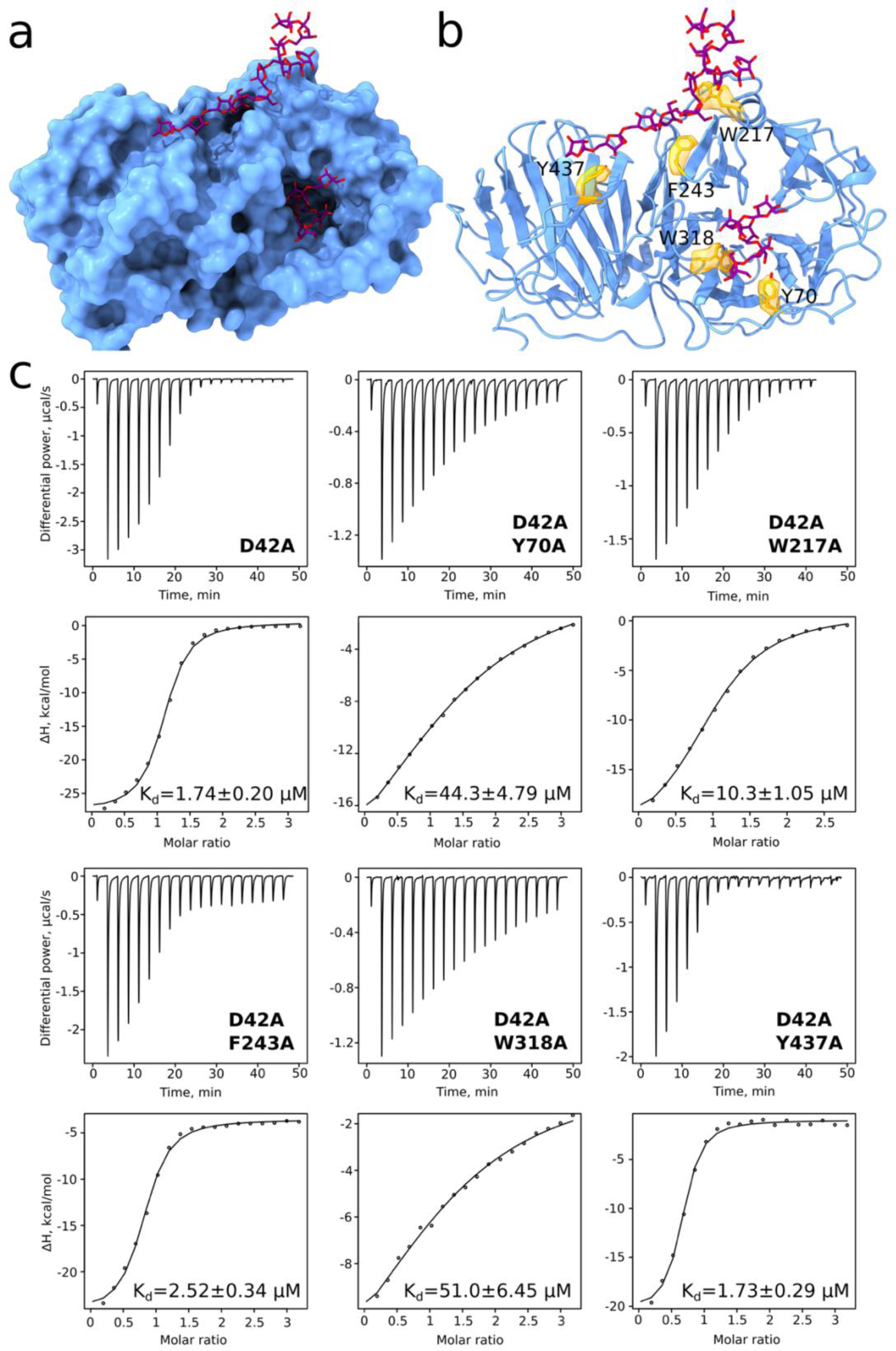
Aromatic residues near the levanase active site, but not the secondary binding site, are responsible for much of the affinity for levan. **a,** Surface representation of the levanase model, with FOS shown as purple sticks. **b**, model and EM density for the residues near the active site (Y70A, W318A) and in the secondary binding site (W217A, F243A, Y437A) that were substituted to alanines are shown in gold. All substitutions were done in the inactive levanase background (D42A). **c**, ITC data from titrations of 8 mg/ml levan into 50 μM of indicated levanase variant. Data fitting assumptions are described in the methods.

**Extended Data Figure 16:**
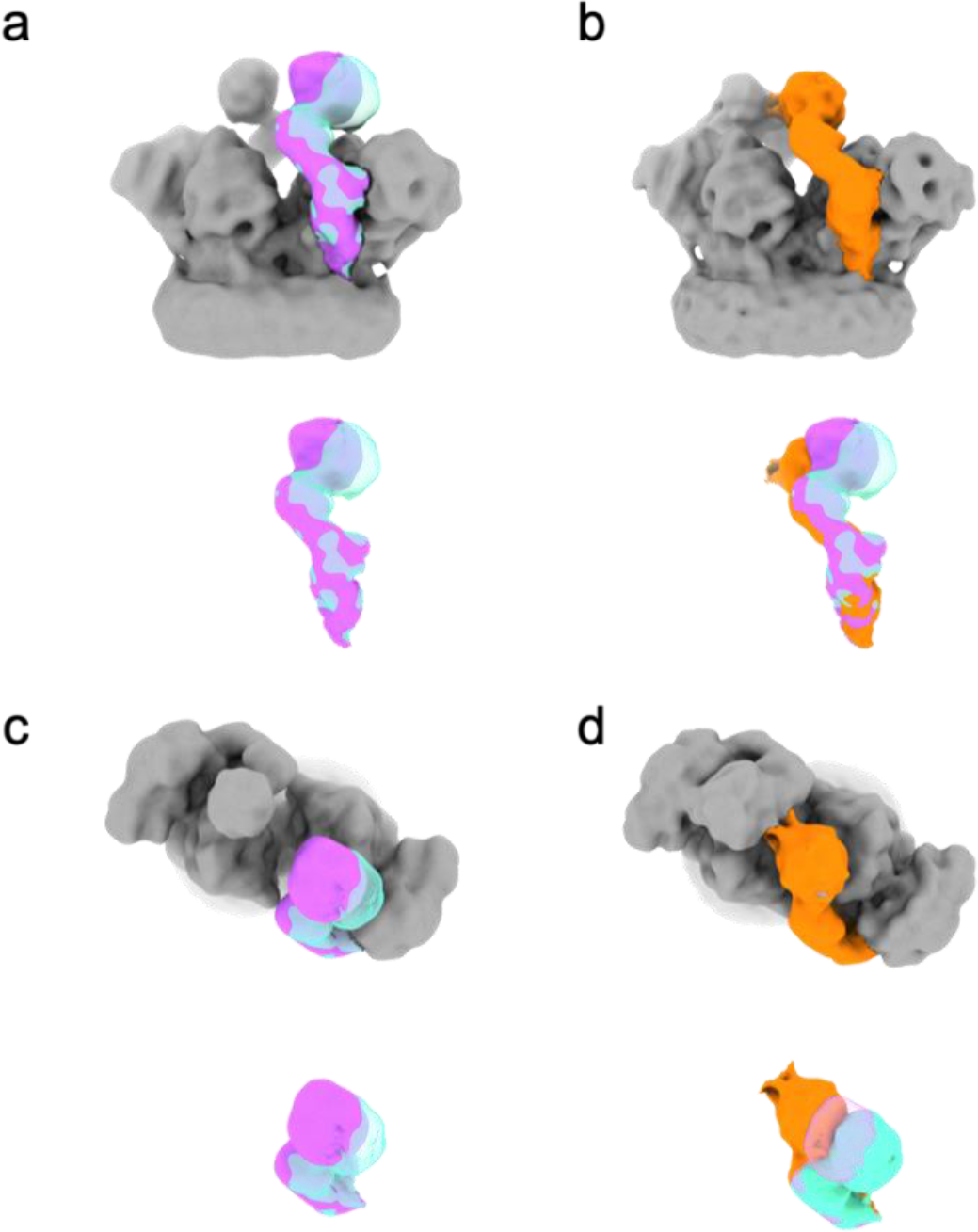
Evidence for cooperative substrate binding by both SGBPs of the levan utilisome. **a**, Variability of the levan SGBP observed for the substrate-bound utilisome with active levanase and short FOS (∼DP8-12). **b**, A novel state observed in 3D classification of the substrate-bound utilisome with inactive levanase and longer FOS (∼DP15-25). Here, one SGBP of the utilisome (orange) appears to reach across and contact the SGBP associated with the other SusC subunit that is present in a docked state. Given that this state was exclusively observed in the presence of long FOS and inactive levanase, we propose that this conformation is the result of both SGBPs in the utilisome interacting with the same stretch of substrate. **c** and **d** show the same maps as **a** and **b** rotated 90° such that they are viewed from the extracellular space.

**Extended Data Figure 17.**
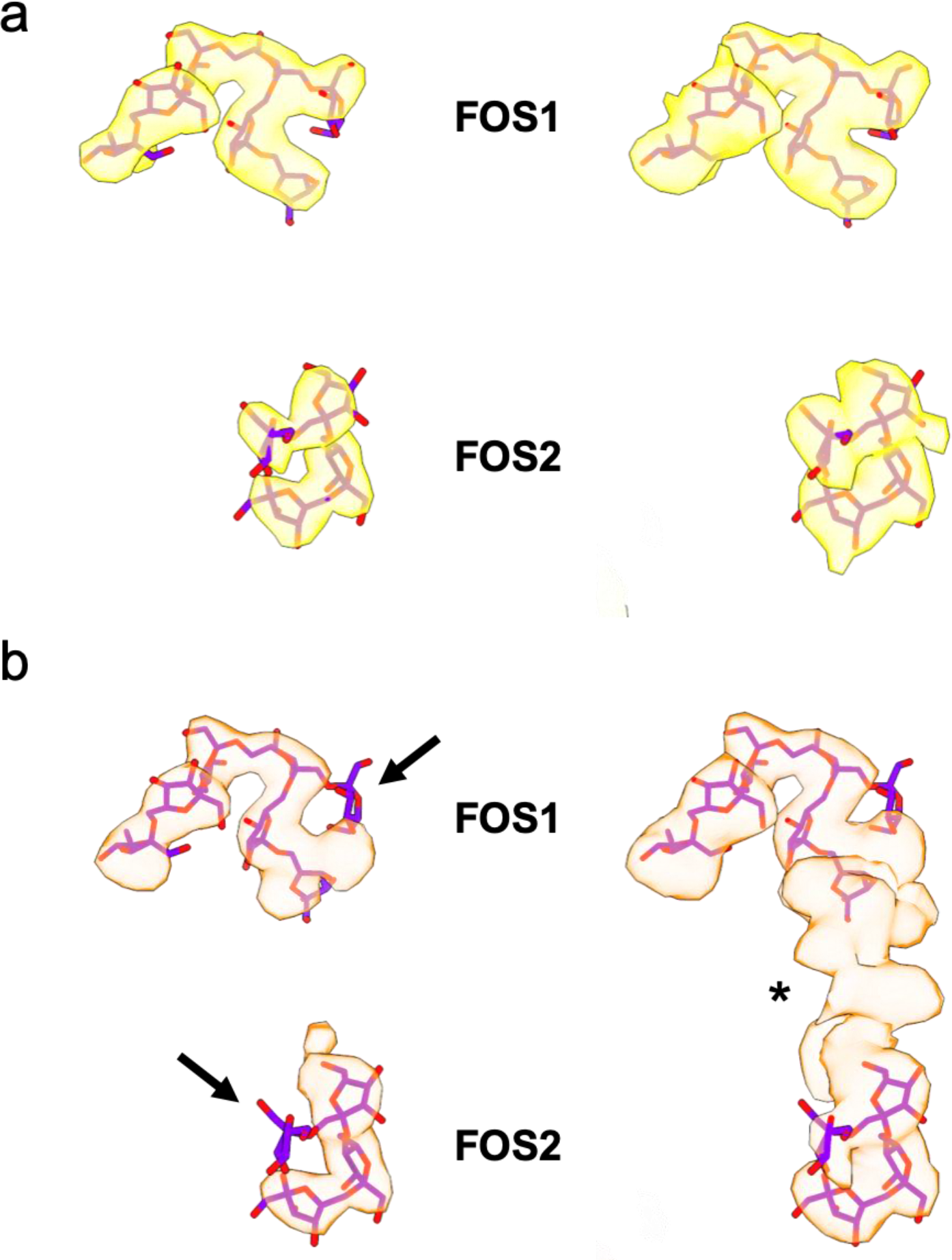
Differences in substrate density for short and long FOS bound in the cavity of SusC. **a**, Isolated FOS density obtained from the levan utilisome dataset with active Bt1760 levanase and short FOS (DP ∼8-12). Density for substrate (yellow) is shown at high (left) and low (right) thresholds. **b**, Isolated FOS density obtained from the dataset with inactive Bt1760 and long FOS (DP ∼15-25). Density for substrate (orange) is shown at high (left) and low (right) thresholds. Arrows indicate missing fructose branches relative to (**a**). Note the presence of unmodeled substrate density bridging the binding sites visible at high thresholds (asterisk). FOS models shown are from the original X-ray crystal structure of the Bt1763-62 SusC_2_D_2_ complex determined in the presence of short FOS (DP6-12)^12^.

